# Connectomic analysis of astrocyte-synapse interactions in the cerebral cortex

**DOI:** 10.1101/2025.02.20.639274

**Authors:** Yagmur Yener, Alessandro Motta, Moritz Helmstaedter

## Abstract

Astrocytes, a main type of glia cells in the cortex, provide metabolic support to neurons, and their possible function as a synaptic partner has given rise to the notion of “tripartite” synapses, suggesting a contribution to neuronal computations. For astrocytes to serve such purposes, the interactions with synapses in neuronal circuits require a level of specificity beyond overall synaptic support. A systematic mapping of the astrocyte-connectome relationship would enable the testing of these hypotheses - such analysis is however still lacking, in particular for circuits in the cerebral cortex. Here, utilizing previously published connectomic data of more than 200,000 synapses, we systematically analyzed the spatial relation between astrocytes and synapses in mouse somatosensory cortex. We developed a quantitative assessment of astrocyte-synapse proximity, finding that only 22.7% of synapses are contacted by astrocytic processes for more than 50% of their synaptic circumference. This non-ubiquitous astrocytic attachment would render astrocyte-synapse specificity plausible. Astrocytic coverage depended strongly on synapse types, with thalamocortical shaft synapses being the most covered by astrocytic processes. We furthermore observed a strong dependence of astrocytic synaptic coverage on synapse size, which was exclusive for excitatory spine synapses. We then investigated the possible relation of astrocytic synaptic coverage to neuronal activity and synaptic plasticity, finding ultrastructural evidence for substantially reduced astrocytic support at synapses consistent with long-term depression, but not for astrocytic coverage dependence on baseline neuronal presynaptic activity. Together, our data demonstrate a high level of specificity of astrocyte-synapse interactions for particular synaptic types. They indicate the potential relevance of astrocytic coverage for synapse stability, in particular for large synapses, suggesting a contribution to long-term maintenance of learned synaptic states. These methods will allow a systematic testing of hypotheses about glial-neuronal interaction in various brain regions, disease models and species including human.

## Introduction

The contribution of astrocytes to the functioning of neuronal circuits has been found to extend from supportive functions (Bergles & Jahr, 1998; Danbolt, 2001; Haydon, 2017; Hertz, 1965; Larsen & MacAulay, 2014; Magistretti et al., 1999; Rothstein et al., 1996; Tanaka et al., 1997) via the formation and elimination of synapses (Chung et al., 2015; Ribot et al., 2021; Ullian et al., 2001) to the suggested involvement in synaptic modulation (Bezzi et al., 2004; Haydon, 2001; Jourdain et al., 2007; Oliet et al., 2001; Parpura et al., 1994; Pascual et al., 2005; Porter & McCarthy, 1996; Schipke et al., 2008), compartmentalized Ca^2+^ elevations potentially selective for the associated synapses (Arizono et al., 2020; Grosche et al., 1999; Kanemaru et al., 2014; Perea & Araque, 2005), and circuit-level computations such as the representation of brain states (Araque et al., 2014; Araque et al., 1999; Cornell-Bell et al., 1990; Deemyad et al., 2018; Ivanov & Michmizos, 2021; Kozachkov et al., 2022; Mu et al., 2019; Perea & Araque, 2005; Poskanzer & Yuste, 2016; Verdugo et al., 2019; Wang et al., 2006)

At the structural level, the relation of astrocytes to synapses has been described as a direct interaction via tripartite synaptic configurations using electron microscopy (EM) (Aten et al., 2022; Calì et al., 2019; Gavrilov et al., 2018; Genoud et al., 2006; Grosche et al., 2002; Grosche et al., 1999; Kikuchi et al., 2020; Lushnikova et al., 2009; Medvedev et al., 2014; Ostroff et al., 2014; Salmon et al., 2023; Špaček, 1985; Tanaka et al., 2013; Thomas et al., 2023; Ventura & Harris, 1999; Witcher et al., 2007). Some of these EM studies revealed that peri-synaptic astrocytic processes are structurally plastic and the level of ensheathment of synapses can be dependent on recent synaptic potentiation (Genoud et al., 2006; Lushnikova et al., 2009; Ostroff et al., 2014; Witcher et al., 2007). Several high-resolution light-microscopy based studies showed the timescale of activity-dependent motility of the peri-synaptic astrocyte microdomains to be minutes (Bernardinelli et al., 2014; Haber et al., 2006) and also possibly dependent on LTP induction (Henneberger et al., 2020).

Several of the suggested astrocytic functions, when analyzed at the level of entire neuronal circuits, require various levels of synaptic specificity of the astrocytic involvement. Yet, a large-scale connectomic analysis of this structural relationship is still missing. Here, we provide such a connectomic analysis of astrocytic synapse associations, allowing us to test predictions about astrocytic function at connectomic scale. We analyze the possible relationship of astrocyte coverage with synapse size, the subtypes of axons and synapses, and determine structural correlates of activity-related astrocytic enlargement. Finally we investigate possible relationships to synapses that may have undergone Hebbian plasticity, or may be in the process of synapse removal.

## Results

### Automated astrocyte reconstruction

We utilized a previously published dense connectomic reconstruction of a 3D-EM volume from layer 4 (L4) of 4-week old mouse barrel cortex ((Motta et al., 2019), Fig. 1a) to obtain a dense reconstruction of astrocytic processes in addition to the previously performed neuronal network reconstruction. We used an automated segmentation of astrocytes (voxelytics, scalable minds, Potsdam, Germany; similar to SegEM (Berning et al., 2015) and TypeEM (Motta et al., 2019)). Astrocytic processes occupied an average of 15.12±1.86% of neuropil volume (mean±s.d. over n=2566 10×10×10 µm^3^ subvolumes sampled from the entire dataset, with nuclei and blood vessels masked-out). The analyzed volume contained 8 astrocyte nuclei and 7 astrocyte somata (Fig. 1b), with one soma containing two nuclei (see example at https://wklink.org/8914), and 89 neuronal cell bodies. Similar to observations in the rodent hippocampus and cortex, astrocytes in our dataset occupied effectively non-overlapping territories (Bushong et al., 2002; Halassa et al., 2007; Ogata & Kosaka, 2002) although synapses positioned near the boundaries of adjacent astrocytes could receive contacts from two astrocytes (Adams et al., 2016; Aten et al., 2022) (Fig. 1c). Each astrocyte territory encompassed tens of thousands of synapses. For subsequent analyses, we integrated these astrocyte reconstructions with pre-existing neurite reconstructions and automatically detected synapses (Fig. 1d).

**Figure 1:**
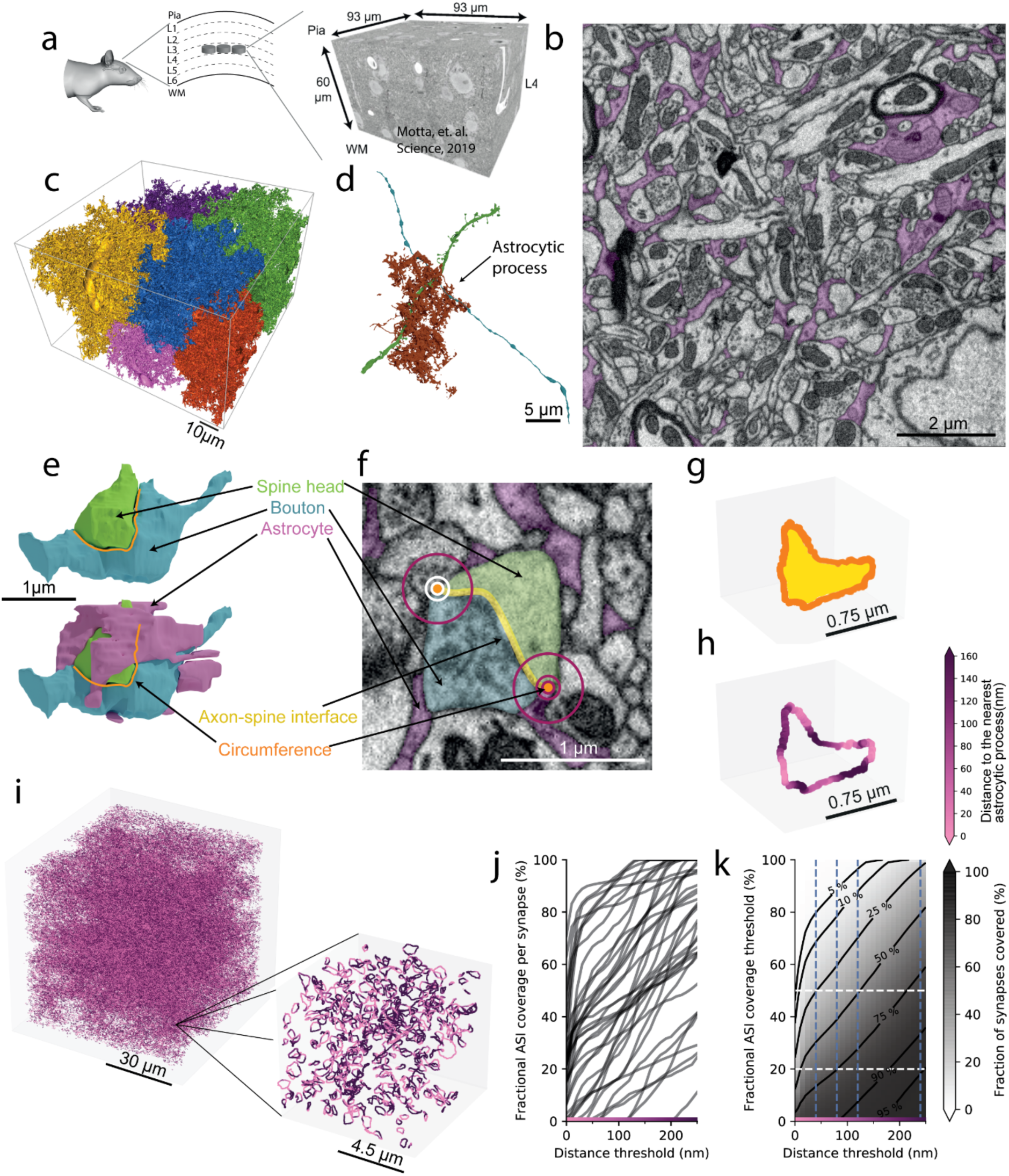
Astrocyte segmentation and quantification of astrocytic synapse coverage. **a,** Sketch of barrels in layer 4 of mouse primary somatosensory cortex (left), 3D EM volume used for all following analyses (60 x 93 x 93 µm^3^, acquired previously including neuropil segmentation, (Motta et al., 2019)). **b,** Automatically detected astrocytic processes (pink) in an example EM cross-section, 2D view. **c,** 3D reconstructions of 7 astrocytes (8 nuclei, 2 share cytoplasm) within this dataset. **d,** An astrocytic process (orange) contacting a synapse with pre-synaptic axon (blue) and post-synaptic dendrite (green). **e,** 3D reconstruction of a spine synapse and an astrocytic process around the synaptic circumference. Top: Pre-synaptic axonal bouton (blue), post-synaptic dendritic spine head (green), and the axon-spine interface (ASI; yellow) and circumference of the synaptic cleft (orange). Bottom: The same synapse with the astrocytic process around the synapse (pink) shown as well. **f,** Cross-section of the same synapse in EM, overlaid labels as in e. For each point of the circumference (orange), the distance to the closest astrocytic process was measured (illustrated for two points of the circumference: circles at 40, 80, 240 nm radius, white if no astrocytic contact, purple if astrocytic contact was detected). **g,h** 3D plot of ASI (yellow), circumference (orange) for one synapse (same as in e,f). Astrocyte proximity to synaptic circumference (h) for each voxel is color-coded by its proximity to the nearest astrocytic surface. **i,** Astrocyte proximity computation applied to (n=227,369) synaptic circumferences in the dataset. Inset, example volume of 9 x 9 x 9 µm^3^ containing n=389 spine synapses, color-coded as in h. **j,** Fractional astrocyte coverage of synaptic circumference in dependence of proximity threshold. For each synapse, fractional coverage is the fraction of voxels that have astrocyte within a distance P_th_, with P_th_ a proximity threshold parameter. Shown for randomly sampled 40 synapses. **k,** Fraction of all synapses that are “covered by astrocytes” quantified in dependence of P_th_ and C_th_, with C_th_ the threshold fraction of a synaptic circumference that was in proximity P_th_ of an astrocyte. Isolines: percentiles. Dashed lines: 20% and 50% synaptic circumference coverage (white), 40, 80 120, 240 nm distance threshold (blue). Approximately 25% of the synapses have at least 50% astrocyte coverage within 40 nm (example synapse: https://webknossos.brain.mpg.de/links/zIicC0ZdwGy8quH8). Approximately 75% of the synapses have at least 20% astrocyte coverage within 80 nm (example synapse: https://webknossos.brain.mpg.de/links/kiRfEw3zg8Fr6NN2).

### Quantification of synaptic astrocyte coverage

To quantify astrocyte – synapse contact we used the following approach: We focused on the synaptic circumference, the zone where synaptic neurotransmitters such as glutamate would diffuse out of the synaptic cleft and could potentially interact with astrocytic processes. This circumference acts as a boundary encircling the synapse (Fig. 1 e-g). For every point along the synaptic circumference, we computed the distance to the nearest astrocytic surface (Fig. 1h), and applied this to 227,369 synapses within the dataset volume (Fig. 1i).

To determine the extent of astrocytic coverage of the synapse circumference, two parameters were relevant: The maximum distance of astrocytic processes to the synaptic circumference that one would still consider as “covering” the synaptic cleft; and the minimum fraction of the synaptic circumference covered at this distance threshold that one would consider sufficient to classify a synapse as “astrocyte-covered”. For any combination of these two parameters, we could then determine the fraction of “astrocyte-covered” synapses (Fig. 1j). At 40 nm distance threshold, the average synaptic circumference “covered” by astrocytes was 31.83±24.42% (mean ± s.d., n=227,369); at 80 nm distance threshold, it increased to 39.26±26.55% (p<10^−300^, Mann-Whitney-U test). When requiring at least 50% of the synaptic circumference to be less than 40 nm from an astrocytic process, only 22.65% of synapses were considered astrocyte-covered; this fraction increased to 77.55% when accepting 20% coverage of each synaptic circumference at 100 nm distance threshold as sufficient (Fig. 1k). Together this implies that, although most of the synapses lay within 100 nm distance to astrocytes, only a small fraction of them can be considered to have substantial astrocyte coverage close-by at their synaptic periphery. For the following analyses, we used 40 nm as the proximity threshold representing the direct contact of astrocyte processes to the synaptic circumference, but controlled the results for sensitivity to the choice of these parameters (Supp. Fig. 7).

### Larger synapses are fractionally more covered

We found a substantial positive correlation between synapse size and the fractional astrocyte coverage (least squares linear fit slope: 24.84% / µm^2^, intercept: 25.61%. Spearman rank correlation coefficient: 0.227, Spearman p-value < 10^−300^, n=154,485, Fig. 2a-c). However, this effect could be attributable to volume displacement: larger synapses occupy larger volume within the neuropil, possibly displacing a larger volume of astrocytic processes towards the synaptic periphery. To control that the size-coverage correlation was in fact a synapse related effect and not a geometric displacement-related one, we applied the same analysis to random contacts between neurites in the densely packed neural tissue (Fig. 2d,e). The size relation to astrocyte coverage was in fact synapse-specific (for non-synaptic interfaces: least squares linear fit slope: 0.52% / µm^2^, intercept: 25.51%. Spearman rank correlation coefficient: 0.105, Spearman p<10^−113^, n= 46,859). Figure 2 e shows the distribution of astrocyte coverage for each separate synapse-size bin. Using this quantification, the effect was even clearer: Each synapse-size bin, on average, exhibited more coverage than the preceding one independently from the others, indicating a gradient of increasing astrocyte coverage with synapse size (p<10^−300^, ANOVA, f-value 547.799; compared to p=0.003 ANOVA, f-value 2.087 for non-synaptic interfaces, with no systematic increase over size bins; see also Supp. Fig. 4d and Supp. Fig. 8a).

**Figure 2:**
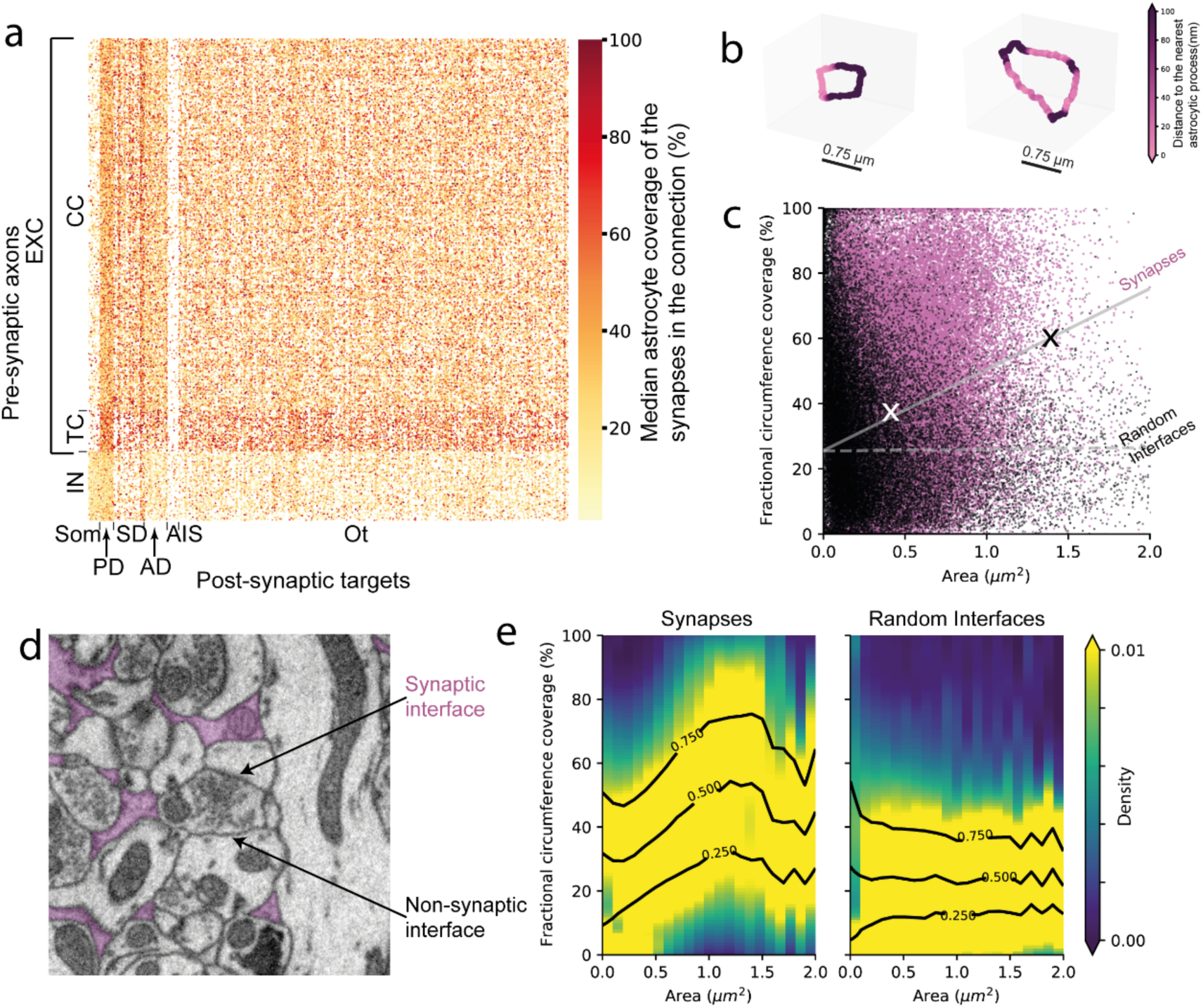
Astrocyte coverage variability in the connectome, and its relation to synapse size. **a,** Astrocyte coverage in relation to the synaptic connectome. Rows: pre-synaptic axons that had minimum output 10 synapses; Columns: post-synaptic targets; dots: synaptic connection between axon and target, color-coded by median astrocyte coverage of the synapses in that connection. **b,** Astrocyte proximity to the voxels of 2 example synaptic circumferences. Synapse on the left is smaller and less covered by astrocytes compared to the larger one on the right. **c,** Relationship between spine synapse size (ASI area) and astrocyte coverage (pink dots, N_syn_=154,485). Larger synaptic interfaces are fractionally more covered by astrocytes (least squares linear fit slope: 24.84% / µm^2^, intercept: 25.61%. Spearman rank correlation coefficient: 0.227, Spearman p-value < 10^−300^). Crosses: example synapses in b. Random neurite interfaces and contact areas as control (black dots, dashed regression line, N_int_=46,859) (least squares linear fit slope: 0.52% / µm^2^, intercept: 25.51%. Spearman rank correlation coefficient: 0.105, Spearman p-value < 10^−113^). **d,** EM slice showing examples of a synaptic interface and a non-synaptic interface within the densely packed neuropil with astrocytes highlighted in pink. **e,** Astrocyte coverage, reported as color map for separate synapse size bins (black lines: coverage isolines). Note substantial dependence on interface size for synaptic (left) but not for random interfaces (right).

### Relation of astrocyte coverage to synapse types

We next analyzed whether astrocytic synapse coverage was related to the types of synapses. First, we measured the astrocyte coverage of shaft synapses, predominantly but not only formed by inhibitory axons, and found no increase in the synaptic circumference coverage with synapse size (Figure 3 a, b) (least squares linear fit slope: -3.11% / µm^2^, intercept: 31.81%. Spearman rank correlation coefficient: -0.063, p<10^−46^, n=51,509. ANOVA between size bins p<10^−83^, f-value 22.951, thus a slight negative slope). Post-hoc tests indicated that one could identify two groups of synapse size bins (Supp. Fig. 8 a). Shaft synapses with smaller contact area (<0.6µm^2^) had significantly more astrocyte coverage (31.84±25.69% mean ± s.d., n=16,145) than the ones with larger contact area (≥0.6µm^2^, 27.90±20.96% mean ± s.d., n=32,971) (p<10^−29^, Mann-Whitney U value: 249,305,864.5). This likely was because of the heterogeneity of the underlying shaft synapse types. Indeed, shaft synapses from excitatory axons onto interneuron dendrites had more astrocyte coverage (median: 39.40%, s.d.: 26.43%, n=8264) and smaller axon-dendrite contact area (median: 0.6 µm^2^, s.d.: 0.46 µm^2^) than the ones from inhibitory axons onto excitatory dendrites (Astrocyte coverage, median: 17.99%, s.d.: 16.95%, p<10^−300^; Contact area, median: 0.96 µm^2^, s.d.: 0.66 µm^2^, n=3874; p<10^−284^; Fig. 3d, Supp. Fig, 8b, c). However, within those subtypes of shaft synapses, there was no synapse size effect on astrocyte coverage (EXC-to-IN Spearman rank correlation coefficient: 0.032, p=0.004; IN-to-EXC Spearman rank correlation coefficient: 0.023, p=0.132).

**Figure 3:**
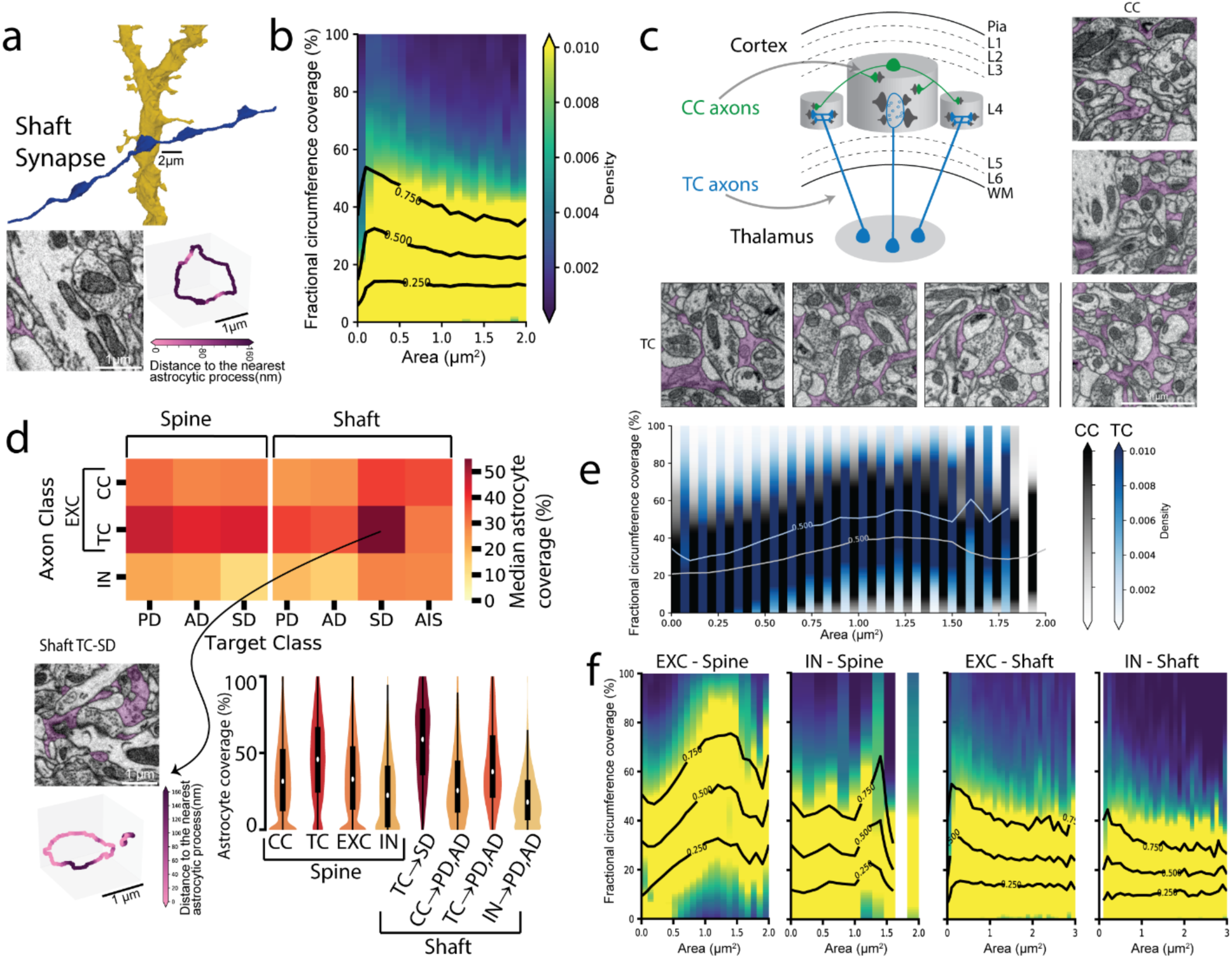
Comparison of astrocyte coverage for different types of synapses. **a,** An example shaft synapse. Top: 3D reconstruction of pre-synaptic axon (blue) and post-synaptic dendrite (yellow). Bottom: EM cross-section (left) and astrocyte proximity to synaptic circumference (right) of this synapse **b,** Astrocyte coverage over synapse size bins showing no synapse size effect on astrocyte coverage for shaft synapses (excitatory and inhibitory combined). Color code and isolines as in Fig. 2e. **c,** Sketch illustrating Cortico-cortical (CC) and Thalamo-cortical (TC) axon types. 3 different examples of each of mono-synaptic CC (right) and multi-synaptic TC (below) axons’ boutons EM cross-sections, with astrocytes highlighted in pink. **d,** Astrocyte coverage of various types of synapses. Top: Median astrocyte coverage of spine and shaft synapses between various axon and dendrite classes. Bottom left: example of an excitatory shaft synapse made by a TC axon onto an interneuron smooth dendrite (SD) covered in astrocytes (pink). Bottom right: Violin plots showing astrocyte coverage distribution of spine and shaft synapses of various connectomic types (example shaft synapses of each connection type: https://webknossos.brain.mpg.de/links/NmWqRntH-PqAAcFs). **e,** Astrocyte coverage over synapse size bins for CC (gray) and TC (blue) spine synapses. TC spine synapses showed more astrocyte coverage than CC. This difference was still visible when accounted for the synapse size effect. **f,** Astrocyte coverage over synapse size bins for excitatory and inhibitory spine and shaft synapses. The relation between astrocytic coverage and synapse size was only present for excitatory spine synapses.

Excitatory spine and shaft synapses had more astrocytic coverage than inhibitory synapses (EXC synapses, median: 31.95%, mean: 34.58%, s.d.: 25.22%, n=138,107. IN synapses, median: 19.80%, mean: 23.46%, s.d.: 19.96%, n=16,637. Mann-Whitney U value: 1,444,782,696.5, p<10^−300^, Fig. 3d,). Synapses of thalamocortical axons were substantially more covered than those of cortico-cortical axons (CC synapses onto spines, median: 31.57%, mean: 33.89%, s.d.: 25.11%, n=95,383. TC synapses onto spines, median: 45.88%, mean: 45.72%, s.d.: 26.32%, n=11,159. Mann-Whitney U value: 393,651,666, p<10^−300^), with TC synapses onto the smooth dendritic shafts of INs showing most coverage (median: 58.91%, s.d.: 26.03%, Kruskal-Wallis H test statistic: 4051.217, p<10^−300^. Bonferroni-adjusted pairwise comparisons with two-sided Mann-Whitney U test p<10^−11^, Fig. 3c, 3d inset).

The increased astrocytic coverage of TC synapses could be caused by the fact that they were overall larger than CC synapses (p<10^−38^, Mann-Whitney U value: 377,126,087.5, n_TC_=9567, n_CC_=85,698), and therefore astrocyte coverage was higher for larger spine synapses. However, even when comparing TC and CC synapses in bins of similar synapse sizes, TC synapses consistently showed more coverage (10^−70^<p<10^−5^, for the most populated synapse size bins; Fig. 3e). Furthermore, the synapse size effect on astrocytic synaptic coverage was still preserved for both classes (Fig. 3e) (CC spine synapses, least squares linear fit slope: 24.58% / µm^2^, intercept: 26.04%, Spearman rank correlation coefficient: 0.231, p<10^−300^, n=95,383. TC spine synapses, least squares linear fit slope: 32.68% / µm^2^, intercept: 34.03% Spearman rank correlation coefficient: 0.326, p<10^−273^, n=11,159).

We concluded that spine synapses were overall more covered than shaft synapses, that the synapse size - astrocyte coverage relation was however only present for excitatory spine synapses, but the largest absolute synaptic coverage was found for TC shaft synapses (Figure 3 f, EXC Spine, least squares linear fit slope: 26.12% / µm^2^, intercept: 26.69%, Spearman rank correlation coefficient: 0.244, p<10^−300^, n=106,542. IN spine, least squares linear fit slope: 2.78% / µm^2^, intercept: 24.80%, Spearman rank correlation coefficient: 0.034, p=0.185, n=1504. EXC Shaft, least squares linear fit slope: -2.21% / µm^2^, intercept: 36.03%, Spearman rank correlation coefficient: -0.046, p<10^−11^, n=22,558. IN Shaft, least squares linear fit slope: -1.03% / µm^2^, intercept: 22.70%, Spearman rank correlation coefficient: 0.004, p=0.713, n=10,545).

### Structural correlates of astrocytic transmitter uptake capacity

Given the astrocytic preference for excitatory spine synapses and their well-established metabolic function in glutamate clearance (Danbolt, 2001; Tanaka et al., 1997), we asked whether we could find structural evidence for astrocytic functional saturation of glutamate clearence. This was of particular interest, since the synaptic size-astrocytic coverage relation found above could be caused by the need for additional astrocytic clearance at synapses that are larger and release more glutamate via their larger synaptic area. If astrocytes operate at saturated uptake capacity, two predictions would follow: first, astrocytic volume fraction should be correlated to synaptic volume density (i.e., the more synapses in a given volume, the more astrocytes are required); and second, synapses along an axon should have more similar astrocytic coverage than synapses from different axons. The latter follows from the assumption that synapses along a given axon are excited by the identical temporal sequences of action potentials, implying that more active axons would release more glutamate than less active ones – a correlation that would be weaker or absent for synapses belonging to different axons.

We first examined the following cases (Fig. 4a): If uptake capacity was not saturated, astrocytes could support multiple synapses without structural changes (Fig. 4a, top row); If their capacity were saturated, in regions of higher synapse density per volume, astrocytes would need to increase their volume density or surface-to-volume ratio to enhance capacity (Fig. 4a, second and third row). Furthermore, when synapses were to get so close to each other as to potentially directly affect each other by glutamate spillover, astrocytes would be predicted to further increase their volume density (Fig. 4a, last row) in order to isolate synapses, reduce synaptic crosstalk and improve the signal-to-noise ratio in neural communication (Tanaka et al., 2013; Witcher et al., 2007).

**Figure 4:**
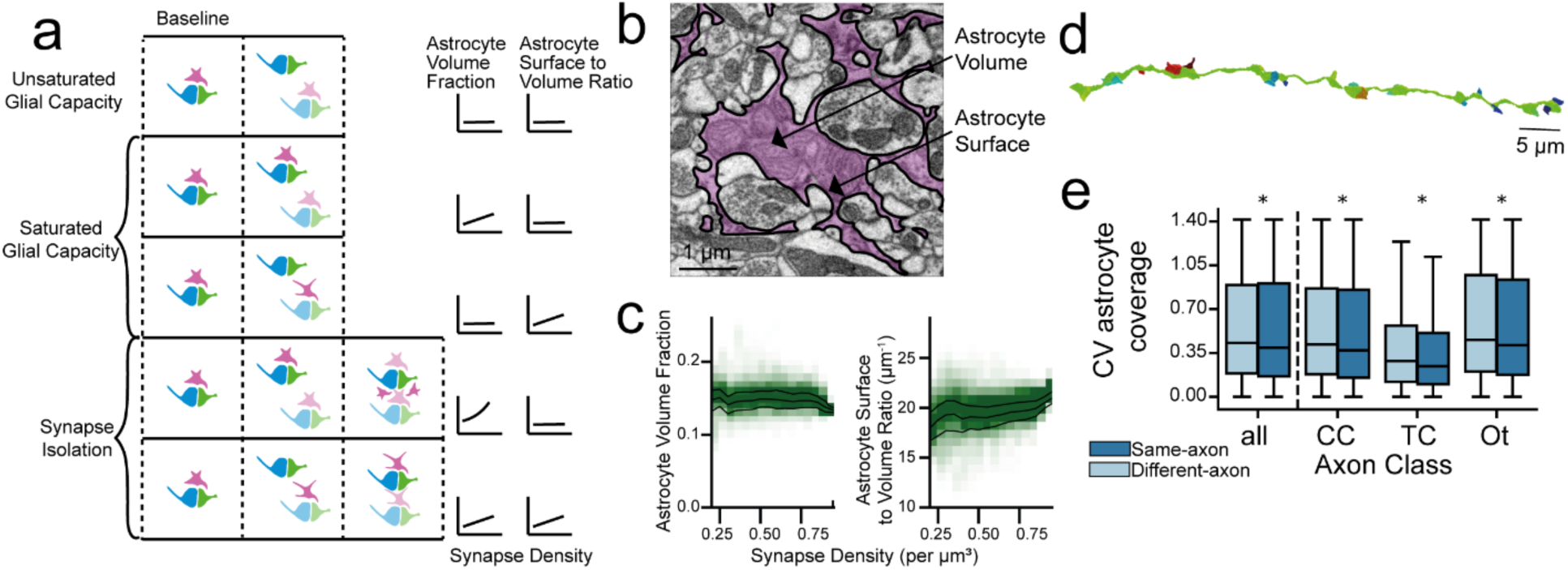
Structural correlates of astrocytic transmitter uptake capacity. **a,** A sketch of possible types of relationship between volumetric synapse density and astrocytic presence. If astrocytic capacity is unsaturated, astrocyte volume fraction is predicted to be independent of volumetric synapse density. If the capacity is saturated, astrocytic volume fraction and/or local surface area is predicted to increase with synapse density. If synaptic isolation is required, astrocytic volume fraction is predicted to show an additional increase beyond a certain synapse density. In all cases, the need for additional astrocytic clearance could be implemented via increased volume fraction or via more complex astrocyte morphology (increased surface-to-volume ratio). **b,** Illustration of surface vs volume of a local astrocytic process: EM slice; astrocyte volume, purple; astrocyte surfaces, solid black lines. **c,** Relationship between astrocytic volume fraction (left) and astrocytic surface-to-volume ratio (right) with the volumetric synapse density (synapses per µm^3^), sampled in 2720 overlapping 10×10×10 µm^3^ boxes. Neither astrocyte volume fraction nor astrocytic surface to volume ratio change substantially in dependence of synapse density, consistent with unsaturated astrocytic capacity (see a, top panel). **d,** 3D reconstruction of an axonal stretch (green) with several post-synaptic spine heads (various colours). The synapses along a given axon are likely to have had a more similar action potential activity history than those located on different axons. **e,** Comparison of astrocytic coverage in synapse pairs from same vs. different axons, with size matching of synapses in both groups (see Methods). Same-axon synapses have significantly more similar astrocyte coverage compared to size-matched different-axon synapses, but the difference is not substantial.

When analyzing the relation of synapse density, astrocyte volume fraction, and astrocyte surface-to-volume ratio, we found no substantial changes in astrocytic density or structure with varying synapse density (Synapse density vs. Astrocyte volume fraction, least squares linear fit slope: -0.004, intercept: 0.155, Spearman rank correlation coefficient: - 0.021, p=0.294. Synapse density vs. Astrocyte surface-to-volume ratio, least squares linear fit slope: 2.055, intercept: 18.758, Spearman rank correlation coefficient: 0.182, p<10^−20^. n=2566, Fig. 4c), in line with unsaturated astrocyte capacity. Astrocyte surface-to-volume ratio was significantly but only slightly higher for higher synapse density regions (<0.7 synapses per µm^3^, 19.85 ± 1.77 µm^−1^, n=501 vs. ≥0.7 synapses per µm^3^, 20.47 ± 1.28 µm^−1^, n=2065; p<10^−15^, Mann-Whitney U value: 396,752). Moreover, average astrocyte coverage of synaptic circumferences did not increase with synapse density (33.55 ± 2.41 % vs. 33.20 ± 1.80 %, p=0.001, Mann-Whitney U value: 468309, Supp. Fig. 3a) Together, the ultrastructural relations do not support a view in which astrocytic clearance capacity is saturated over the range of observed synapse densities, and no indication of synaptic isolation at very high synapse volume densities.

### Same-axon variability of astrocytic coverage

If the variability of astrocytic coverage of synapses were driven by the electrical activity (and thus synaptic release frequency) of presynaptic axons, we would predict synapses of the same presynaptic axon to be more similar in astrocytic coverage than those from different axons. We therefore analyzed the astrocytic coverage of synapses made by the same presynaptic axon and compared it to the astrocytic coverage of synapses sampled from different presynaptic axons (Fig. 4e). For this, we had to additionally control for synapse size (which we had found to impact astrocytic coverage, see Fig. 2). We matched different-axon synapse pairs to the synapse size of the same-axon pairs for comparison. We found that the astrocyte coverage of same-axon synapses was in fact more similar to each other than the astrocyte coverage of different-axon synapses even when accounted for their synapse size (Fig. 4 e). This effect was statistically significant, and would be consistent with astrocyte motility towards more active synapses (Genoud et al., 2006; Haber et al., 2006; Lushnikova et al., 2009) (Same-axon synapses CV, median: 0.39, mean: 0.57, s.d.: 0.49, n=5000. Different-axon synapses CV, median: 0.43, mean: 0.58, s.d.: 0.48, n=15,000. Mann-Whitney U statistic: 36,059,972, p<10^−4^). However, the difference was not substantial, in line with the above conclusion that glial capacity of astrocytes was unsaturated, and thus the requirement for astrocytic recruitment to more active synapses was modest.

### Relation of astrocytic structure to possible Hebbian synaptic plasticity

We finally explored astrocytic coverage in the context of same-axon-same-dendrite (SASD) synaptic pairs, which have been used to analyze a possible history of Hebbian synaptic plasticity, in particular LTP and LTD ((Bartol et al., 2015; Dorkenwald et al., 2022; Motta et al., 2019; Sievers et al., 2024), Fig. 5a). These SASD analyses had identified synaptic pairs with low relative size variance and large synaptic size (consistent with LTP, Fig. 5a, top), and synaptic pairs with low relative size variance and small synaptic size (consistent with certain forms of LTD, Fig. 5a, bottom), leaving the remaining synaptic pairs with mixed synapse sizes as neither consistent with saturated LTP nor LTD (based on a set of additional assumptions, see Methods).

**Figure 5:**
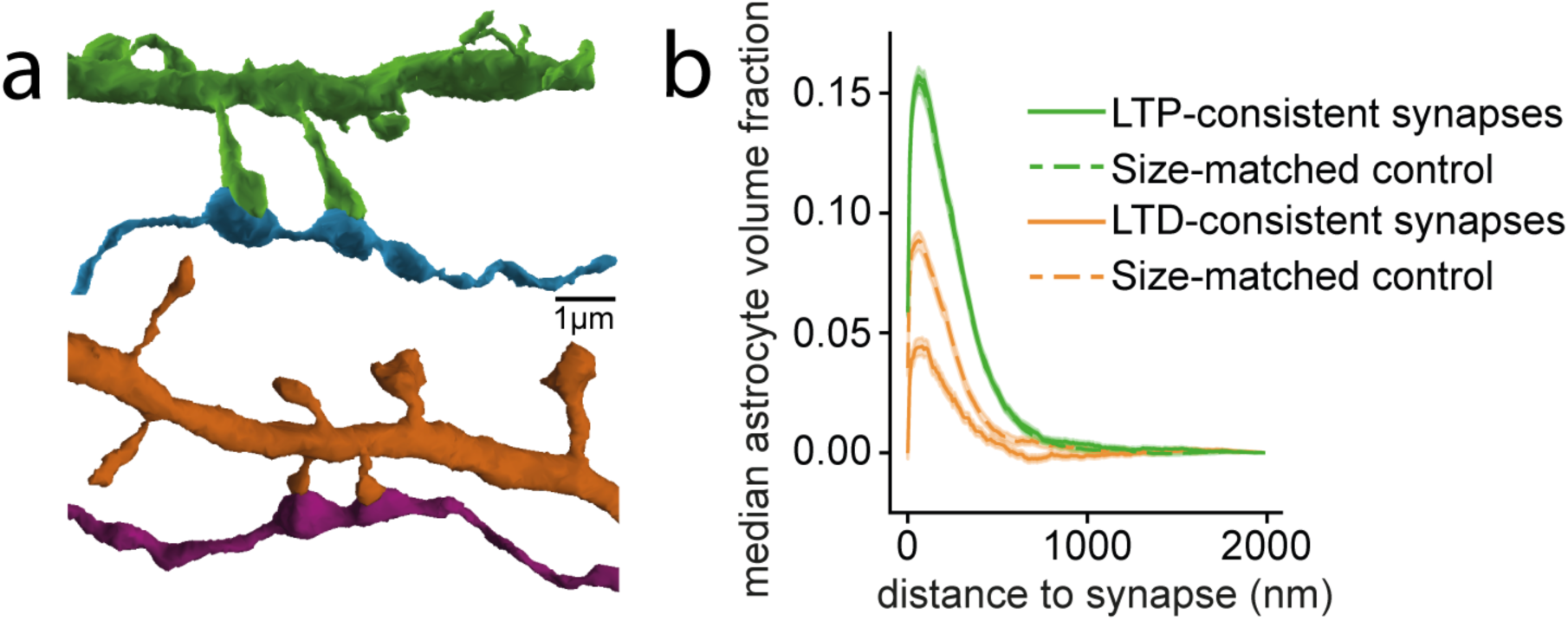
Relation of astrocytic structure to possible history of Hebbian synaptic plasticity. **a,** Example 3D reconstructions of large (top) and small (bottom) same-axon same-dendrite (SASD) synapse pairs. Within each pair, the synapses are of similar size, consistent with a history of saturated Hebbian plasticity, see Motta et al., 2019. (Dendrites (green, orange), axons (blue, magenta)) **b,** Median astrocyte volume fraction over distance to the synapse, shown for synapses from SASD pairs consistent with a history of LTP (solid green), and consistent with a history of LTD (solid orange). Dashed lines: size-matched controls from synapses not associated with LTP- or LTD-consistent SASD synapse pairs. Note that LTD-consistent, but not LTP-consistent synapses have almost 2-fold reduced astrocytic coverage compared to the size-matched synapses without evidence for LTD, suggesting an effect of astrocytic coverage on spine synapse stability (Mann-Whitney-U test p<10^−9^ for LTD-consistent synapses, p=0.64 for LTP-consistent synapses).

We wondered whether, beyond the size effect on astrocyte coverage, we could detect differences in astrocyte coverage between SASD synaptic pairs consistent with LTP/LTD, and those that were not consistent with this form of plasticity. For the analysis of astrocytic coverage, we first matched the size distribution of synapses found in non-Hebbian consistent pairs to each Hebbian-consistent category through subsampling. We found that large and overly similar (LTP-consistent) synapses, as expected due to their size, had more astrocytic volume fraction than small and overly similar (LTD-consistent) synapses. However, when comparing the astrocytic coverage of large and overly similar (LTP-consistent) synapses to that of size-matched non-SASD synapses, we found no significant difference (Astrocyte volume fraction above baseline within 100 nm of the synapses. LTP-consistent, median: 15.00%, mean: 16.12%, s.d.: 16.81%, n=2038; Non-SASD size-matched controls, median: 15.37%, mean: 16.42%, s.d.: 17.15%, n=2038. Mann-Whitney U statistic: 2,058,972, p:0.637).

When repeating the same analysis for small and overly similar (LTD-consistent) synapses, to our surprise, LTD-consistent synapses had almost half the amount of proximal astrocytic volume fraction compared to size-matched non-SASD synapses (Fig. 5b, astrocyte volume fraction above baseline within 100 nm of the synapses. LTD-consistent, median: 4.21%, mean: 6.49%, s.d.: 16.09%, n=1938. Non-SASD sized-matched control, median: 8.02%, mean: 9.98%, s.d.: 17.40%, n=1938. Mann-Whitney U statistic: 1,663,742, p<10^−9^).

LTD-consistent synapses also had less astrocyte coverage than their sized-matched controls non-Hebbian synapses (original: 20.23%, sampled: 28.33% of the synapses had more than half of their synaptic periphery covered by astrocytes).

The substantial differences in astrocytic coverage and volume would be in line with a model in which synaptic strength reduction via LTD may involve a retraction of astrocytic structural support from spine synapses, which would in turn underscore a function of astrocytes for spine synapse stability.

## Discussion

We used large-scale 3D EM data of mammalian cortex and its connectomic reconstruction to examine hypotheses about glia-neuronal interaction using structural observables. We found that (1) Only a minority of synapses are substantially covered by astrocytes, when quantified as at least 50% close contact; (2) The relative extent of astrocyte ensheathment was positively correlated to synapse size; this was not explained by glial volume displacement at larger synapses; (3) Astrocytic coverage of synapses was strongly dependent on synapse type, with spine synapses covered more than shaft synapses for each target type, with the exception of thalamocortical shaft synapses onto interneuron dendrites standing out as the most strongly covered synapses overall. (4) The relation of relative astrocyte ensheathment to synaptic size was exclusive to excitatory synapses onto dendritic spines, suggesting a unique astrocyte-synaptic interaction at these synapses; (5) Astrocytic ensheathment did only weakly depend on structural indicators of increased or correlated synaptic activity, suggesting astrocytic neurotransmitter uptake capacity to be unsaturated; (6) Astrocyte-synapse coverage was notably reduced at synaptic pairs that were consistent with ongoing synaptic depression (synapse size reduction). Together, our data points to a particular association of astrocytes with large spine synapses, and suggests these interactions primarily provide structural synaptic stability rather than enhanced homeostatic capacity.

### Tissue preparation effects

Chemical fixation is known to suppress extracellular space (ECS, (Van Harreveld et al., 1965)) by as much as 85% (Korogod et al., 2015). This could increase the observed apparent astrocytic coverage of synapses (Korogod et al., 2015; Ohno et al., 2007). The rather low amount of astrocyte-covered synapses in our data (Figure 1 k) is therefore still an overestimate. Since we were primarily interested in synaptic coverage comparisons between synapses within the same dataset, we did not correct for the ECS shrinkage effect.

### Synapse size estimation by contact area

We use the area of the axon-target interface as a proxy for synapse size, drawing on previous findings that link axon-spine interface (ASI) area and spine head volume to synaptic strength indicators such as the post-synaptic density (PSD) area, the active zone (AZ) area, and vesicle count in the synaptic bouton (Bopp et al., 2017; Cheetham et al., 2012; Harris & Stevens, 1989; Schikorski & Stevens, 1997) and in line with previous connectomic studies (Bartol et al., 2015; de Vivo et al., 2017; Motta et al., 2019). For shaft synapses, however, the relationship between axon-dendrite contact area and PSD area is less explored. While the lack of relationship between shaft synapse size and astrocytic coverage in our data is striking, there could be more subtle relationships to synaptic strength of shaft synapses that get masked by the rather coarse measure of shaft synapse interface area and the lack of ECS. Studies relating morphological features of shaft synapses to shaft synapse strength will allow more detailed analyses in the future.

### Previous analyses of astrocytic synaptic coverage

For quantifying the amount of astrocytic synaptic coverage, two main parameters have to be set a-priori: (1) What is the maximum distance between a synapse and an astrocytic process such that this configuration constitutes “coverage” (and what structure is the measurement relating to: synaptic cleft center, synaptic cleft circumference, presynaptic bouton, postsynaptic spine head); and given a proximity distance definition: (2) what fraction of a given synapse’s circumference should be associated with an astrocytic process for this synapse to be considered “covered”. Once these parameters have been set, an overall fraction of synapses with astrocytic coverage in a given tissue can be determined. The choice of these parameters varies widely in the literature; yet, when accepting these various choices, the resulting fraction of synapses with astrocytic coverage has been reported to be 25-45% in most studies (Ostroff et al., 2014; Špaček, 1985; Ventura & Harris, 1999; Witcher et al., 2007), with the exception of ferret cortex (1.4-24.4% coverage (Thomas et al., 2023)) and cerebellum (75-100% coverage, (Grosche et al., 2002; Lippman et al., 2008; Špaček, 1985)). Our data would be consistent with the 25-45% coverage range.

However, the broad range of sometimes implicitly chosen parameters, reported across many species, brain regions and ages (40-90% of synapses are covered (Aten et al., 2022; Genoud et al., 2006; Kikuchi et al., 2020; Lushnikova et al., 2009; Ostroff et al., 2014; Rollenhagen et al., 2015; Špaček, 1985; Tanaka et al., 2013; Thomas et al., 2023; Ventura & Harris, 1999; Witcher et al., 2007)), makes comparisons difficult. Our attempt at quantitative assessment, and the opportunity to analyze results with any chosen set of distance and coverage parameters (Figure 1 j, k) was aimed at allowing a systematic analysis of astrocyte-synapse association. Applying the same approach to other brain samples will provide a systematic comparative assessment of the variability (and key determinants) of synapse-astrocyte association.

### Previous analyses of synapse size and astrocytic synaptic coverage

Previous studies have repeatedly shown that synapses that have astrocyte contact were larger than those without (Kikuchi et al., 2020; Thomas et al., 2023; Witcher et al., 2007) and larger synapses with more complex PSD shapes were more likely to have astrocytic contact (Lushnikova et al., 2009; Ventura & Harris, 1999; Witcher et al., 2007). However, astrocyte contact length at the synaptic perimeter was not proportional to the synapse size in rat hippocampus and amygdala (Calì et al., 2019; Gavrilov et al., 2018; Ostroff et al., 2014; Ventura & Harris, 1999; Witcher et al., 2007), with the exception of (Arizono et al., 2020; Herde et al., 2020; Lushnikova et al., 2009). All of these studies agree that in hippocampus there was either an inverse or no correlation between synapse size and the fraction of the perimeter covered by astrocytes. In contrast to hippocampus, the results are more consistent in cortex where synapses with larger PSDs were contacted by larger astrocytic processes (Salmon et al., 2023) and synapse size positively correlated with astrocytic perimeter length (Genoud et al., 2006; Thomas et al., 2023). The latter work in L2/3 of ferret visual cortex, however, showed no correlation between synapse size and fraction of periphery covered by astrocyte (Thomas et al., 2023). Our data, showing strong relationship between synapse size and *fractional* astrocytic coverage for excitatory spine synapses suggests the relative coverage to be an important readout. Future work will have to determine whether using this analysis approach, synaptic astrocytic coverage fractions are more consistent across model systems and brain regions.

### Connectomic type selectivity of astrocytes

Astrocytes have been reported to more likely contact asymmetric synapses than symmetric synapses (Kikuchi et al., 2020) and cover spine synapses more than shaft synapses (Gavrilov et al., 2018), and within the shaft synapse population asymmetric shaft synapses have more frequent astrocytic contact than symmetric shaft synapses (Ostroff et al., 2014). Our data is in line with astrocyte coverage fractions being higher for excitatory over inhibitory and spine over shaft synapses, but a detailed analysis of synapse type relationship revealed the most covered cortical synapses to be excitatory shaft synapses from TC axons (Fig. d) – indicating that astrocytic synaptic coverage has strong subtype specificity and thus needs to be analyzed at detailed connectomic level.

### Relationship of astrocyte-synapse contact to presynaptic activity

Astrocytic plasticity has been reported to be associated with pre-synaptic glutamate release and astrocytic mGluR activation (Bernardinelli et al., 2014; Perez-Alvarez et al., 2014) or to also require the involvement of post-synaptic NMDARs (Henneberger et al., 2020; Lushnikova et al., 2009). Following whisker stimulation, an increased expression of astrocytic glutamate transporters without a significant increase in the total volume or surface area of astrocytes has been reported (Genoud et al., 2006). Moreover, under baseline conditions, (Salmon et al., 2023) report that astrocytic mitochondria are positioned closer to denser clusters of synapses. Together, these findings may indicate that adaptation of astrocytic processes to increased demand of glutamate clearance is primarily achieved by non-structural means, in line with our finding of a lack of relation of astrocytic volume fraction or surface-to-volume ratio to local synapse density. This is also supported by our data showing an only modest effect on astrocytic coverage of same-axon versus different-axon synaptic pairs (Fig. 4 d, e). By comparing same-axon synapse pairs to size-matched different-axon synapse pairs, we controlled for the substantial bouton size heterogeneity and dynamics that occur even within one axon (Chéreau et al., 2017; Sammons et al., 2018; Stettler et al., 2006).

Our additional finding of existence of the strong synapse size-to astrocytic coverage correlation (Fig. 2) on the other hand suggests that the structural impact of synapse size on increased astrocyte coverage may be more closely associated with synaptic stability than with metabolic needs.

### Astrocytic structural plasticity and synapse stability

Time-lapse light microscopy and EM analyses have reported that astrocytic processes exhibit spontaneous motility (Bernardinelli et al., 2014; Haber et al., 2006; Hirrlinger et al., 2004) which can be enhanced via stimulation (Bernardinelli et al., 2014; Perez-Alvarez et al., 2014), resulting in increase of both the percentage of synapses contacted by astrocytes and the extent of their coverage (Genoud et al., 2006; Lushnikova et al., 2009; Wenzel et al., 1991). Some reports have however contradicted these findings (Henneberger et al., 2020; Ostroff et al., 2014). Larger and more stable spines have furthermore been shown to have stable contact with astrocytic process that are less motile (Arizono et al., 2020; Bernardinelli et al., 2014; Haber et al., 2006). These data are consistent with our results that large spine synapse size alone is sufficient to explain substantial astrocytic coverage irrespective of it being consistent with LTP (Fig. 5).

In contrast, our finding that small spine synapses consistent with a history of LTD show *lower* astrocytic coverage than the non-LTD-consistent sized-matched controls may imply that loss of astrocytic support is associated with loss of synaptic stability.

Overall, our data indicate that astrocyte-synapse relations may serve primarily the goal of synapse stability, and astrocytic coverage variability may not be dominated by metabolic or homeostatic requirements linked to synaptic activity.

### Outlook

With our approach of quantitative analysis of astrocyte-synapse associations at the connectomic scale (involving hundred of thousands of synapses), a method for screening astrocytic properties across cortices, brain regions, age, species and pathological conditions has been put forward, which will provide a substantial foundation for studies of glial function and glial contribution to synaptic network operation in our and other animals’ brains.

## Acknowledgements

We thank Kristen Harris and Mark Ellisman for enlightening discussions and Sahil Loomba for feedback on an earlier version of the manuscript, Christian Guggenberger at MPCDF Garching for excellent HPC support, Daniel Werner and the scalableminds team for running the latest version of astrocyte type predictions and segmentation, and Heiko Wissler for visualizations.

## Author Contributions

MH conceived, initiated and supervised the project; YY developed and conducted all analyses with input from MH and AM; AM developed an earlier version of the astrocyte detection networks; YY and MH wrote the paper with input from AM.

## Funding

All research was funded by the Max Planck Society.

## Methods

### 3D EM dataset

The dataset analyzed in the present study was published previously (Motta et al., 2019). All procedures regarding animal experiments, tissue preparation, and dataset acquisition were described there. The dataset has previously been used in (Berning et al., 2015; Boergens et al., 2017; Motta et al., 2019; Schmidt et al., 2024; Staffler et al., 2017). The dataset was sized 8274×5338×3321 voxels at a voxel size of 11.24×11.24×28 nm³, yielding a total size of (92.6×61.8×94.8) µm³.

### Connectomic type definitions

Synapse detection, neurite reconstruction, and neurite type definitions were conducted as described in (Motta et al., 2019): Briefly, synapses were automatically detected using SynEM (Staffler et al., 2017). Excitatory and inhibitory axons were defined by their spine or shaft synapse preference rates, respectively. Excitatory axons had more than 50% spine synapse fraction and inhibitory axons had less than 20% spine synapse fraction, with the remaining axons defined in the “other” class (N_exc_=5894, N_inh_=893, N_oth_=528). Within the excitatory axon class, Thalamocortical axons were identified based on the work of (Bopp et al., 2017) where it was observed that the individual boutons of thalamocortical axons in S1 usually formed multiple synapses, which allowed reliable identification in EM without specific labelling. Post-synaptic targets such as somata, spiny dendrites of excitatory cells, smooth dendrites of INs, axon-initial-segment, and apical dendrites of neurons located in deeper cortical layers were identified using semiautomated heuristics to detect these subcellular compartments.

### Automated Detection of Astrocytes

To automatically classify astrocytes, we used the Voxelytics software (scalable minds GmbH, Potsdam, Germany, developed in collaboration with MPI Brain Research, Dept. of Connectomics), similar to the dense reconstruction method described in (Loomba et al., 2022). Neurite reconstruction and automated synapse detection were performed as previously reported in (Motta et al., 2019).

First, a classifier was trained to assign type probabilities to each voxel in the dataset to belong to one of the following classes: astrocyte, dendrite, axon, spine head, and other. Then, another set of classifiers were trained to assign voxel affinities to each voxel (i.e. the connectivity to neighboring voxel pairs) (Lee et al., 2017), followed by watershed and hierarchical agglomeration to obtain object segmentations (see (Motta et al., 2019) for details). Then these two results were combined: voxel-wise type probabilities were averaged over the voxels in a segmentation object. In addition, the type probabilities per object were used in the hierarchical agglomeration process as follows: the astrocyte, dendrite, and axon classes were normalized (with the spine head class included in the dendrite class). Then, objects were classified as astrocytes if the astrocyte type score was larger than 0.4 and was the maximum of all type scores. Lastly, all voxels belonging to an object classified as astrocyte were classified as astrocyte.

To evaluate astrocyte classification accuracy, we first used manually volume-annotated astrocytic processes in two bounding boxes each 2×2×2µm^3^ in size using webknossos (Boergens et al., 2017). To focus on the performance near synapses, we then annotated astrocytes near 30 randomly sampled synapses. For the second case, the ROI for evaluation was obtained by 6-fold iterative binary dilation of the segment objects belonging to the synapse (with a 3×3×3 voxel^3^ object, yielding 67 ..193 nm surrounds). Together, voxel-wise precision and recall were 0.87, and 0.95. We furthermore qualitatively assessed peri-synaptic astrocytic processes as well as thicker astrocytic processes by visual inspection.

### 3D visualization of separate astrocytes

To visualize separate astrocytes contained in the dataset, first, astrocyte soma coordinates were obtained using an earlier version of the dataset segmentation and type prediction (similar to (Motta, 2016)) as follows. Connected segments were identified by applying connected components on a segment-edge list which yielded a list of separate agglomerates, each made up of many segments. Then, agglomerates that had minimum 500 segments whose average neurite score was lower than 0.75 and average astrocyte score was larger than 0.5 were identified which yielded 8 separate large astrocytic processes. To find the soma location of astrocytes, first, all the nuclei in the dataset were identified by selecting agglomerates that had minimum 0.6 nuclei score. The subset of nuclei that had the largest contact area with astrocytic processes were identified as astrocyte nuclei, the centroids of these nuclei were registered as astrocyte soma coordinates. Upon visual inspection of the EM data, one of the identified large astrocyte processes did not have a soma as it was a branch of an astrocyte that was outside of and neighboring to the dataset. For the remaining 7 large astrocytic processes, 8 nuclei were found.

To visualize the 3D reconstructions of these 7 astrocytes, the Voxelytics and Webknossos demo by Scalable Minds GmbH based on (Motta et al., 2019) publicly available at https://wklink.org/8914 was utilized. Using the soma coordinates of 7 astrocytes, the precomputed isosurface meshes was loaded in Webknossos and downloaded each as separate STL files. There were 6 meshes as two of the astrocyte agglomerates had merged during the new version of the segmentation process. Then, we re-generated the high resolution renderings of the 3D reconstructions using Amira Software (Stalling et al., 2005).

### Definition of synaptic circumference

To define the synaptic circumference for each synapse, we proceeded as follows. For each automatically detected synapse, a bounding box of 4×4×4 µm^3^ centered at the synaptic center position was defined. The segmentation objects of the presynaptic and postsynaptic neurites and any astrocytes present in the bounding box were upsampled to 10 × 10 × 10 nm^3^ voxel size to obtain an isotropic representation. Then, the axon-spine interface (ASI) for spine synapses (or the synaptic interface for all other synapse types) was defined as the collection of voxels that were neighbors of both the pre- and postsynaptic segments. Then, in order to define the voxels of the circumference of the synaptic interface, we applied multiple steps of 3D mathematical morphological operations implemented using SciPy (Virtanen et al., 2020). The following operations use a binary structuring element of size 3×3×3 voxel^3^ of square connectivity equal to one for two iterations. First, we applied binary dilation to each of the synaptic compartments and obtained their overlapping voxels resulting in a flat 3D object containing the synaptic interface. Then, binary closing operation was applied to both of the synaptic compartments at once, merging them at the synaptic interface. We then subtracted the closed object from the dilated-overlap object. The resulting binary mask was taken to be the synaptic circumference for this synapse. (Fig 1 g, h).

### Astrocyte proximity to synaptic circumference

In order to determine the distance of astrocytes to the voxels of a synaptic circumference, all astrocyte segments within the bounding box were converted to a binarized mask, and the distance transform to this mask was computed. Then, for each voxel of the synaptic circumference, the value of the distance transform was recorded as astrocyte proximity. These computations were also performed in the up-sampled data representation at 10×10×10nm^3^ voxel size.

### Astrocyte coverage of synaptic circumference

For a given synapse, to compute the astrocytic coverage of its synaptic circumference, the astrocyte proximity was thresholded at a given proximity threshold *P_th_* and the fraction of circumference voxels with astrocytic proximity below *P_th_* was recorded as coverage fraction for this synapse. This was applied to all synapses in the dataset.

### Fraction of synapses covered by astrocytes

To obtain the fraction of synapses in sufficient proximity to astrocytes to be considered *covered* by astrocytes (a necessary condition for the concept of “tripartite” synapses), the minimum astrocytic coverage of a synapse’s periphery was used as a parameter (threshold of astrocyte coverage fraction per synapse, *C_th_*; see Fig. 1 j, k for the effect of *P_th_* and *C_th_* on synaptic coverage results). For all following analyses, *P_th_* was set to 40 nm. To control for the sensitivity of the results to the particular choice of parameters, all analyses were also conducted for the parameter choices *P_th_* =20..250 nm with step size 10 nm and 250..600 nm with step size 50 nm (Supp. Figs. 7-9), yielding similar results.

### Area of the axon-spine or neurite interface

To determine the area of synaptic and non-synaptic interfaces, we first obtained the collection of interface voxels (at 10nm upsampled resolution) as described above (“Definition of synaptic circumference”). We then approximated the collection of interface voxels as a cylindrical object of height 60 nm, used the marching cubes algorithm (Lorensen & Cline, 1987b) to determine this object’s approximate surface area, and calculated the cylinder cross section area as 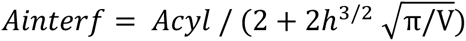, with *V* the total volume of the interface voxels and *Acyl* the surface area of the collection of interface voxels. *Ainterf* was then interpreted as the synaptic or non-synaptic interface area. The cylinder height, *h*, (at 60 nm) was obtained from comparison of *Ainterf* for axon-spine interfaces to the methods in Motta et al 2019.

### Manual inspection of synapse size effect on astrocyte coverage

To sample from synapses of various size homogeneously, we randomly selected 3 synapses each from 10 size bins (spaced 0.1 µm^2^, range 0.05 µm^2^ – 1.5 µm^2^). For each of the 30 sampled synapses, a bounding box of 2×2×2µm^3^ was centered around the synapse and the astrocyte voxels were manually annotated using the volume tracing mode in Webknossos. Astrocyte coverage was computed using the above described method. The synapse size - astrocyte coverage relation was also found (Supp. Fig. 2 c).

### Stacked kernel density visualization of synapse size-coverage relation

To obtain a visualization of the relationship between synapse size and astrocytic coverage that would allow a judgement of the size effect for each synaptic size range independently, we proceeded as follows. First, synapse sizes were binned in 0.1 µm^2^ increments. For each bin, a kernel density estimation of the astrocyte coverage distribution for the synapses in that bin was computed. The kernel density estimates for each size bin were then stacked along the x-axis, normalized, and the 25^th^, 50^th^ and 75^th^ percentiles were plotted as lines (Fig. 2e; 3b,e,f) .

### Relation of astrocyte coverage to interface size for non-synaptic interfaces

To control for tissue geometry (in particular astrocytic volume displacement at large synapses) in the synapse-size-to-astrocyte-coverage relation, we randomly sampled 48,779 agglomerate contacts (i.e. axo-axonic, dendro-dendritic, and axo-dendritic contacts) from the automated neurite reconstructions and computed interface area and astrocyte coverage as described above for synaptic contacts.

### Computation of synapse density, astrocyte volume fraction and surface-to-volume ratio

To analyze the relation of astrocytes to local synaptic density, we iterated over the segmented volume of the L4 dataset with a 10×10×10µm^3^ bounding box with 5×5×5µm^3^ step size. At each iteration, we computed the spine synapse density, total astrocyte volume (total astrocyte voxel volume), total astrocyte surface area (surface area estimated via marching cubes algorithm (Lorensen & Cline, 1987a), average spine synapse area, total spine synapse area, total spine head volume, and the average astrocyte coverage of individual spine synapses. This resulted in 2720 (10×17×16 iteration in x-y-z) bounding box measurements. These measurements were confined to the neuropil volume by masking out nuclei, blood vessels, and large dendritic bodies within a bounding box. For the synapse density analysis, we included only those bounding boxes where at least half of the volume consisted of neuropil n=2566.

### Analysis of astrocyte coverage for same-axon synapses

To analyze the variability of astrocytic synaptic coverage for pairs of synapses from the same axon versus pairs of synapses from different axons, we considered only non-inhibitory axons with at least 2 spine synapses. For this analysis, the proximity threshold for astrocytic synaptic coverage was set to 120 nm, yielding for a more symmetric astrocytic coverage distribution across synapses. Similar results were obtained for other proximity thresholds. To control for the strong synapse size effect on astrocytic synaptic coverage (see Fig. 2), it was necessary to ensure that the synapse pairs from different axons had a similar size distribution as those from same-axon synapse pairs. This was implemented as follows:

For each axon class, n=5,000 same-axon synapse pairs and size matched different-axon synapse pairs were randomly sampled. For this, first, 2 randomly sampled spine synapses s1,s2 were sampled from a randomly selected axon. Then, all spine synapses from all other axons were binned according to their size (in 0.1 µm^2^ bins). Then, a pair of synapses s1’ and s2’ were randomly sampled from the size bins that s1 and s2 were from, while ensuring that s1’ and s2’ were from different axons. Then, for each of the synapses s1, s2, s1’, s2’ the astrocytic coverage fraction *fcovastro* was computed. Finally, for each synapse pair (s1,s2), (s1’,s2’), (s1,s2’),(s1’,s2), the coefficient of variation (C.V.) of the synapse’s astrocytic coverage was computed as *cvastro* = *sqrt*(2) ∗ *delta*(*fcovastro*)/ (*sum*(*fcovastro*) + *epsilon*), with *epsilon* = 0.0001 to avoid division by 0.

As a result, we obtained two C.V. distributions per axon class: the distribution of C.V. of astrocytic coverage for same-axon synapse pairs (H1: (s1, s2)) and for size matched different-axon synapse pairs (H0).

For this analysis, all axons were intentionally split at their branch points to minimize agglomeration merger errors (similar to (Motta et al., 2019)). This also reduced the possible effect of action potential failure at axonal branch points, which would confound the interpretation of same-axon synapse pairs as having been exposed to similar electrical (action potential) activity.

### Astrocytic volume fraction in dependence of distance to synapse

We measured the astrocyte volume fraction in dependence of the distance to a synapse as follows. For each spine synapse, we computed the distance transform from the synaptic interface voxels. Then, we measured the astrocyte volume fraction in ranges of increasing distance from the synaptic interface (as determined by the distance transform for distances of 10 nm .. 2000 nm at 10 nm increment). For the computation of astrocyte volume fraction, segmentation objects belonging to nuclei, blood vessels, the presynaptic axon and the postsynaptic spine head were masked out. This yielded for each synapse a distance profile of astrocytic volume fraction. The distance profiles were smoothed using a moving average window of 20 nm with step size 10 nm. The profiles were aligned by subtraction of the baseline astrocyte volume fraction, which was taken as the astrocyte volume fraction in the outermost shell for each synapse. We then computed the astrocyte volume profile for each synapse by multiplying the baseline-subtracted astrocyte volume fraction per shell with the respective shell volume (corrected for masked voxels, see above), and summing over all shells.

### Analysis of astrocytic coverage of same-axon same-dendrite synapse pairs

Previous work (Motta et al., 2019) had identified a significant overabundance of same-axon same-dendrite (SASD) synapse pairs with above-random synapse size similarity, which occurred for large and small synapse sizes, respectively. Here, we used these previously identified classifications of SASD synapse pairs as large&similar, small&similar, and remainder (interpreted in the context of Hebbian plasticity, see Motta et al 2019 and Suppl. Fig. 6a). We compared the astrocyte volume fraction for these SASD synapse pair classifications. In particular, we asked whether synapses that were a member of large&similar pairs had a different astrocytic neighborhood than synapses of similar size that were part of the remainder pairs. Similarly, we asked whether synapses that were a member of small&similar pairs had a different astrocytic neighborhood than synapses of similar size that were part of the remainder pairs. For this, we first sampled a set of single synapses from the remainder pairs for which the distribution of synapse sizes matched the shape of the distribution of single synapse sizes of the large&similar distribution (note that for this analysis, we considered the synapses of SASD pairs as single synapses, with however the association to the respective SASD paired synapse pools as class labels), and analogously for the small&similar synapse size distribution (Suppl. Fig. 6 a). We only considered synapses from remainder pairs with c.v. of their synapse sizes of at least 0.3 to ensure non-similarity.

Size matching of the synapse distributions was achieved by random sampling (with replacement) of single synapses from the large&similar pool, for which a sized-matched synapse from the remainder pool was randomly drawn (sizes were matched within bins of 0.07 µm^2^ width, range 0.21-1.25 µm^2^). Analogously, a size-matched pool of remainder synapses was drawn for the small&similar synapses (sizes were matched within bins of 0.012 µm^2^ width, range 0.046-0.220 µm^2^).

Then, the astrocytic neighborhood of the synapses in the respective pools were analyzed (Fig. 6). The comparison of astrocyte volume fraction above baseline within 100 nm of the synapses for different synapse pools was reported but the result was consistent within 40 nm to 500 nm.

### Statistical methods

The following statistical tests were performed and are documented in the code repository at https://gitlab.mpcdf.mpg.de/connectomics/astrocyte/-/tree/astro?ref_type=heads.

For synapse size-astrocyte coverage relationship of spine, shaft synapses as well as the control experiment of random interfaces and connectomic subtypes, a Spearman rank correlation coefficient was reported with a Spearman p value (Figures 1 c-e; 2 b, f). Additionally, ANOVA was applied for comparison of synapse size bins followed by post-hoc Tukey’s HSD test (Supp. Fig. 8a).

For astrocyte coverage distribution comparison of EXC and IN axons’ synapses, a one-sided Mann-Whitney U test was applied. For multiple comparisons of axon and target subtypes in the violin plot (Figure 3 d) with TC axons’ shaft synapses onto smooth dendrites, both a Kruskal-Wallis H test and a Bonferroni-adjusted pairwise comparisons with two-sided Mann-Whitney U test were applied and the highest p value was reported. For comparison of both the synapse size distributions and the astrocyte coverage distributions of all CC and TC axons’ synapses, one-sided Mann-Whitney U test was applied (Figure 3 c, d). Additionally, for pairwise comparisons of astrocyte coverage distributions of CC and TC axons’ synapses for each size bin, two-sided Mann-Whitney U test was applied with Bonferroni correction (by multiplying resulting p by number of size bins) (Figure 3 e). For comparison of synapse size bins in EXC, IN, spine, and shaft synapse astrocyte coverage plots, Spearman rank correlation coefficient was reported with a Spearman p value (Figure 3 f).

For analysis of structural correlates of astrocyte capacity looking at relations between synapse density, astrocyte volume, and astrocyte surface to volume ratio (Figure 4 c); a Spearman rank correlation coefficient was reported with a Spearman p value. Additionally, for comparison of higher and lower synapse density regions one-sided Mann-Whitney U test was applied.

For comparison of astrocyte coverage of same-axon synapse pairs to different axon partner-size-matched synapse pairs (Figure 4 e), a one-sided Mann-Whitney U test was applied.

For comparison of the distributions of astrocyte volume fraction within 100 nm of different classes of same-axon same-dendrite synapse pairs (Figure 5), a one-sided Mann-Whitney U test was applied.

## Supplementary Figures

**Supp. Fig. 1.**
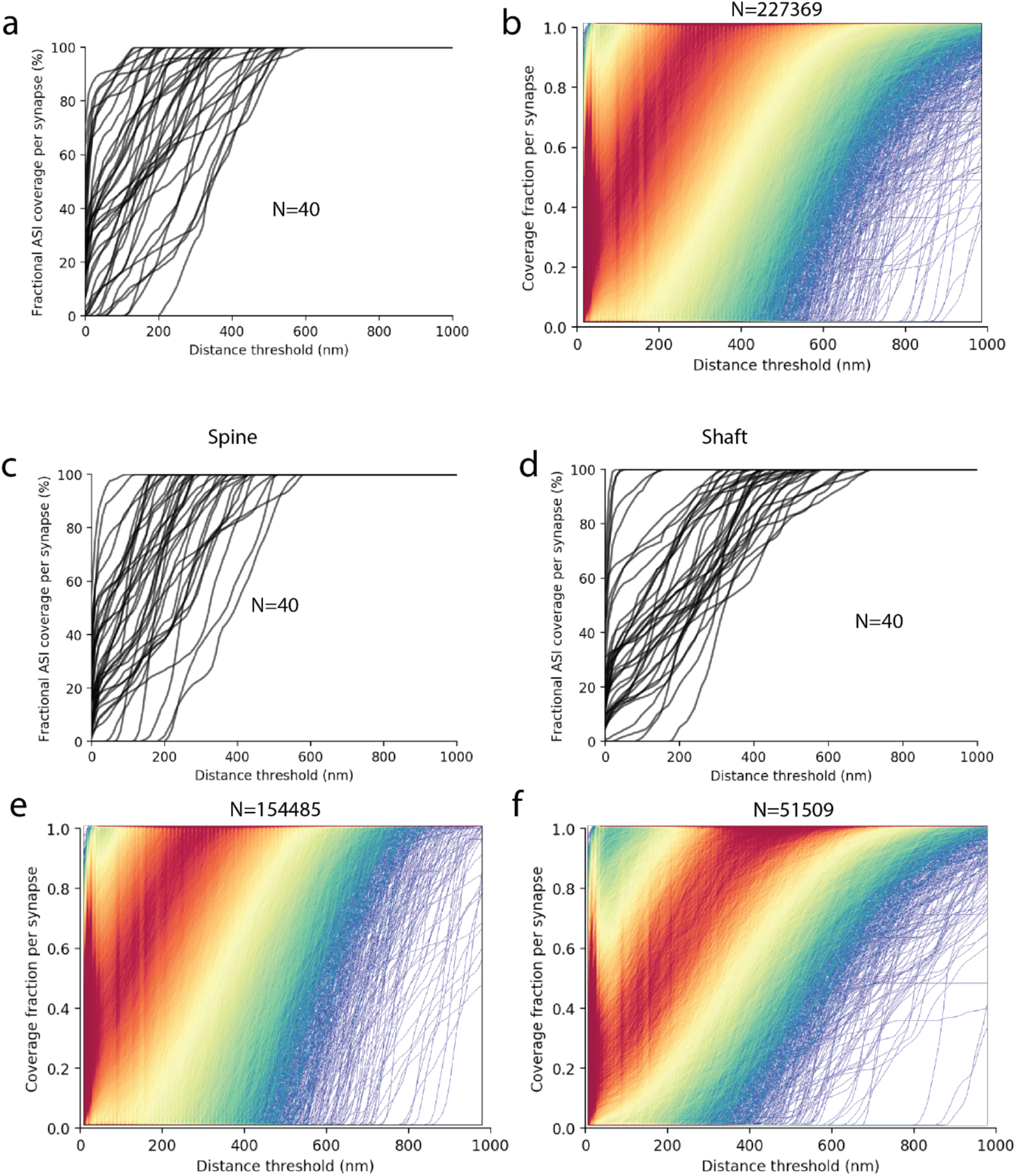
**a,** Fractional astrocytic coverage of the ASI circumference, in dependence of the maximum distance at which the synapse was considered “covered”, shown for n=40 example synapses. **b,** Similar to a but with all n=227,369 synapses shown. Color indicates curve density. **c,** As in a shown for spine synapses, only. **d,** As in a shown for shaft synapses, only. **e,f** As in b, restricted to spine (e) or shaft (f) synapses.

**Supp. Fig. 2.**
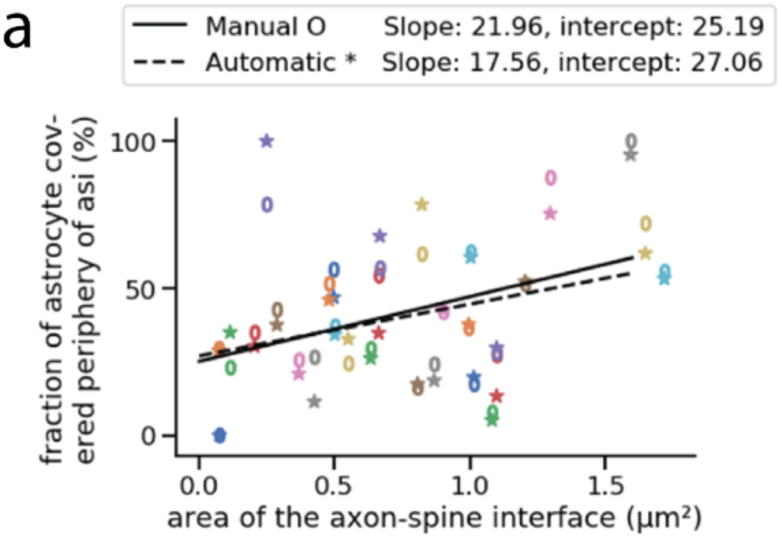
Validation of synapse-size to astrocytic coverage relation from manually annotated astrocytes for 30 randomly sampled synapses from various synapse size bins.

**Supp. Fig. 3.**
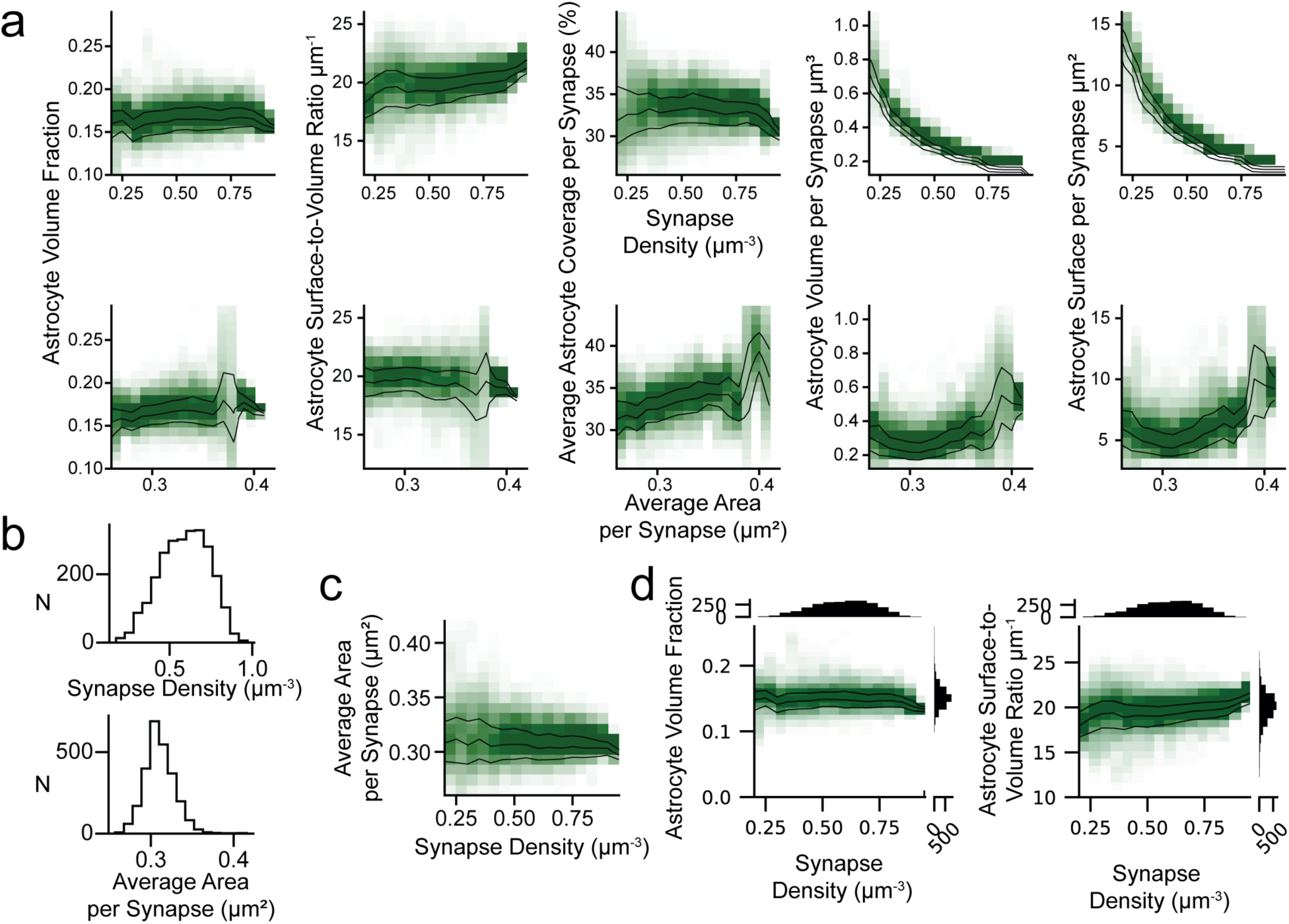
Relationship of astrocytic volume fraction to volumetric synaptic variability (supplementary to Fig. 4). **a,** Astrocytic properties in dependence of synapse density and synaptic area per synapse. None of the additional astrocytic properties substantially increased with synapse density. Average astrocyte coverage, astrocyte volume and surface per synapse only increased with average synapse area (Average synapse area vs average astrocyte coverage, Spearman rank correlation coefficient: 0.283, p<10^−44^, n=2566). **b,** Variability of synapse density and average synapse area within the bounding boxes. **c,** Synapse density vs. average area per synapse (Spearman rank correlation coefficient: -0.069, p=0.000487, n=2566). **d,** Same as in Fig 4c but with histograms as insets.

**Supp. Fig. 4.**
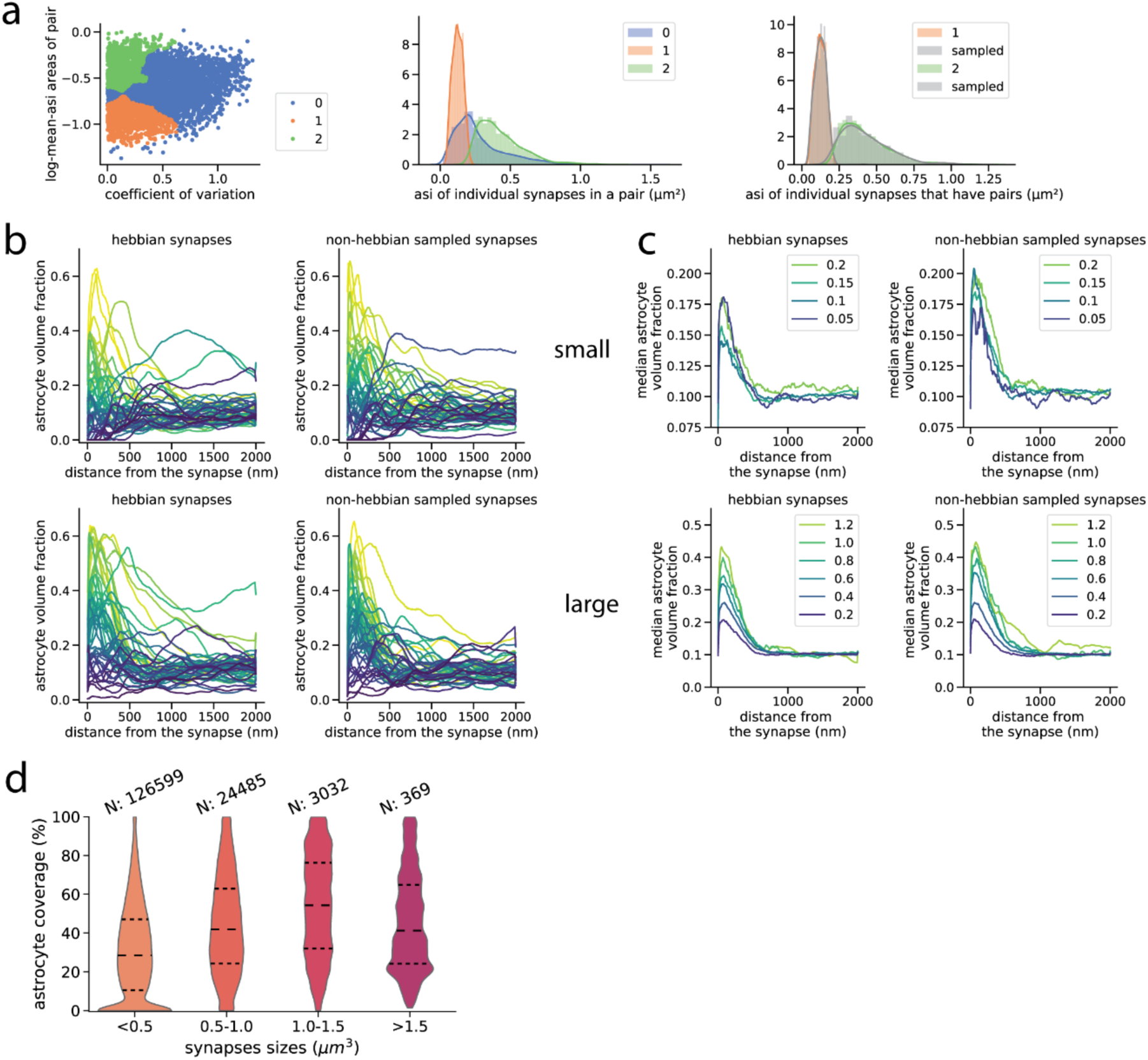
Supplementary to Fig. 5. **a,** Left: The SASD synapse pairs previously classified as non-Hebbian (blue), LTD (orange) and LTP (green) classes (Motta et. al., 2019). Middle: the distributions of synapse sizes for each class. Right: The distribution of synapse sizes after non-Hebbian synapse size distribution (previously blue, now grey) was resampled to match each Hebbian synapse classes’ size distribution. **b,** Astrocyte volume fraction over distance from synapse for example synapses from different classes. Top: small synapses. Bottom: Large synapses. Left: Hebbian synapses. Right: non-Hebbian sampled synapses. **c,** Median astrocyte volume fraction over distance from synapse for synapses in different size bins. Top: small synapses. Bottom: Large synapses. Left: Hebbian synapses. Right: non-Hebbian sampled synapses. **d,** (Henneberger et al., 2020), suggest that large size-saturated synapses might be uncovered by astrocytes to allow spillover to induce new synapse formation. Here we show all primary spine synapses divided into 4 size bins. Indeed, the largest of synapses have a drop in astrocyte coverage. Note that, only 0.2% of the synapses fall into this size bin.

**Supp. Fig. 5.**
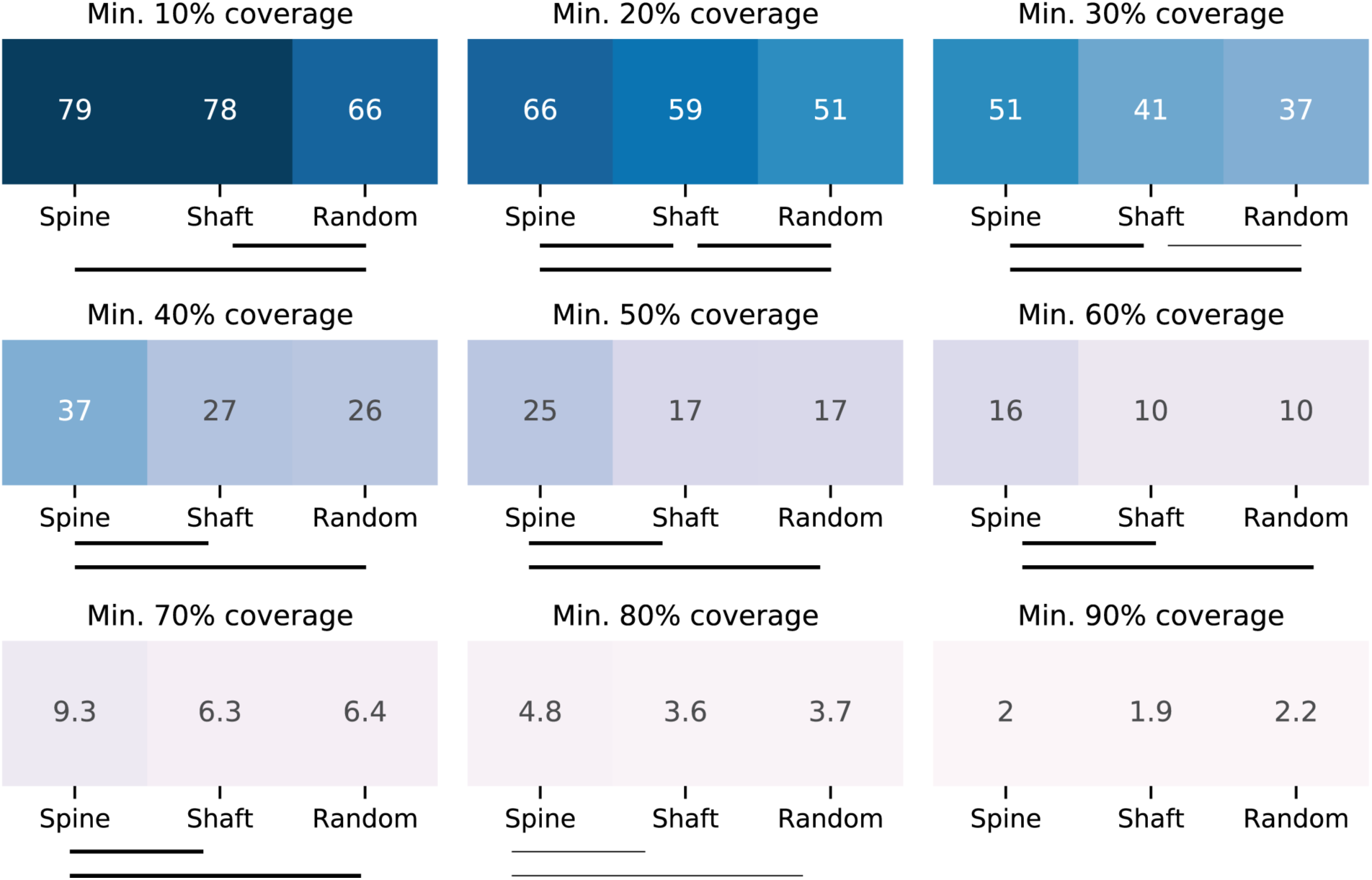
Readout of fraction of synapses covered, in dependence of the circumference coverage threshold. Bars and bar thickness indicate significance (p>10^−6^ no bar, P<10^−93^ thick bar) two proportion z test converted to p values with Bonferonni multiple comparison adjustment (x3) N_spine_=154,485, N_Shaft_=51,509, N_rand_=46,859

**Supp. Fig. 6.**
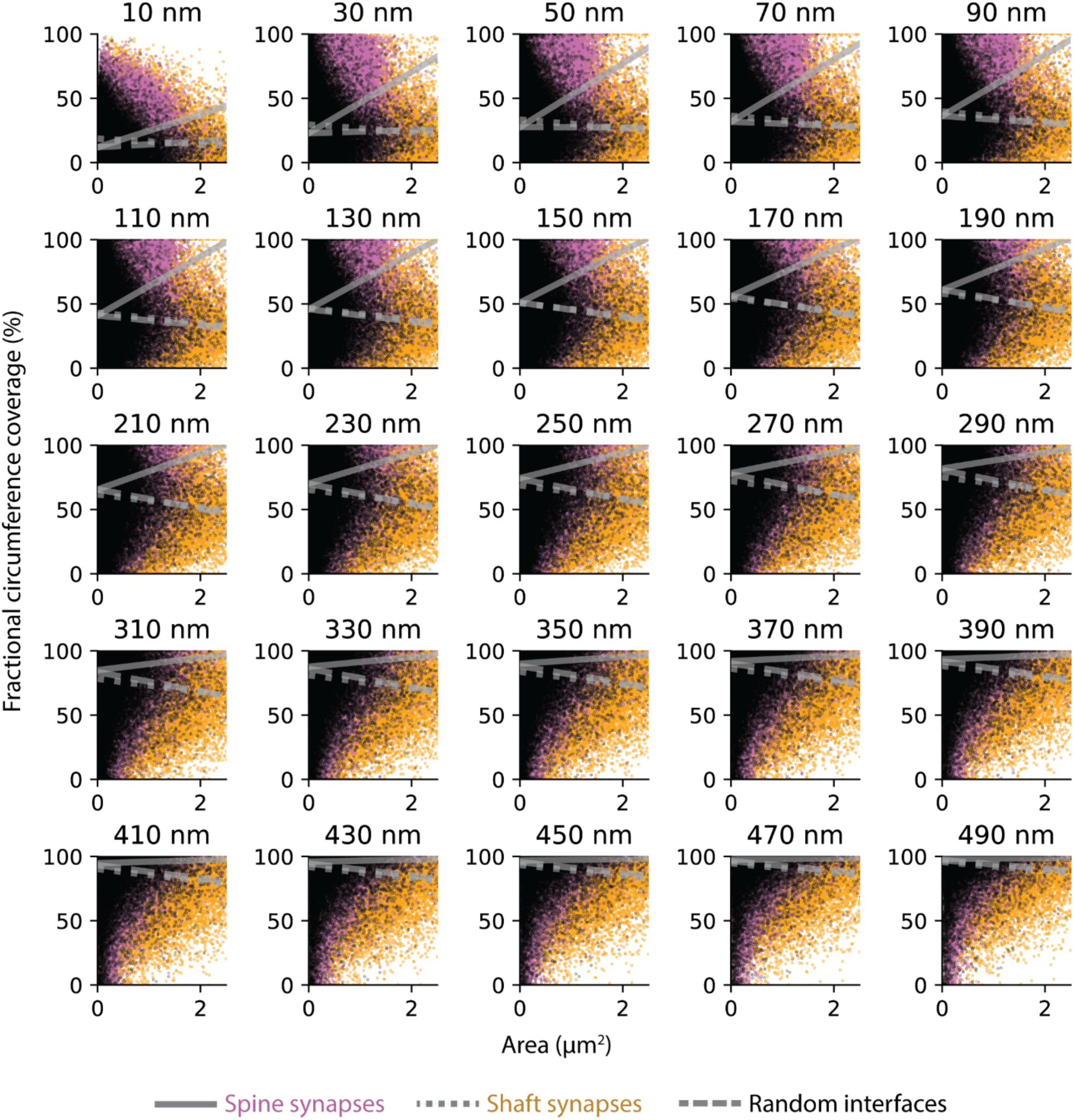
Dependence of astrocytic coverage on synapse size, reported for spine and shaft synapses and random interfaces in dependence of various distance threshold values in range between 10 – 500 nm at 20 nm step size.

**Supp. Fig. 7.**
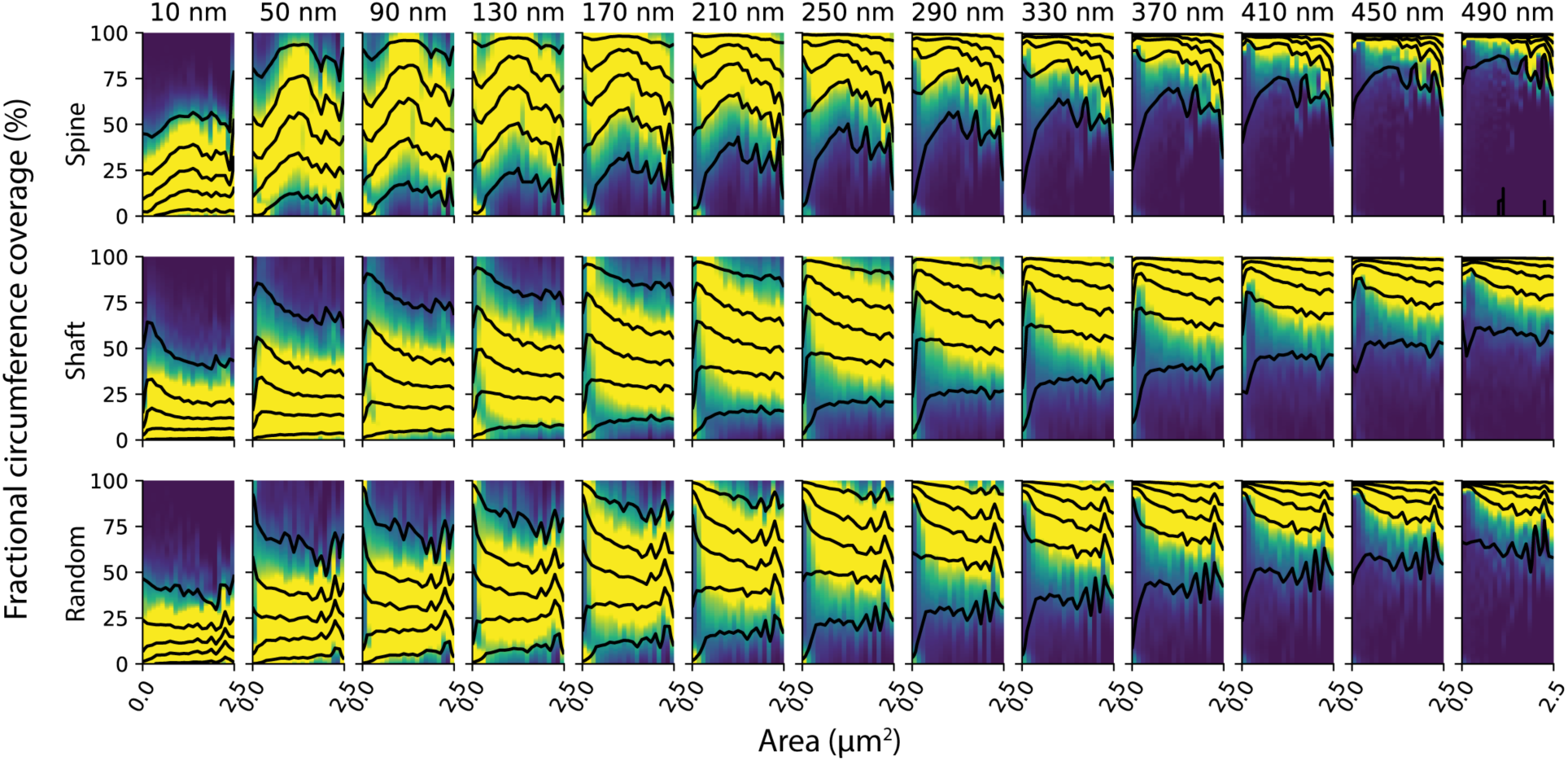
Stacked kernel density visualization of synaptic and nonsynaptic area relation to fractional astrocyte circumference coverage for spine and shaft synapses and random interfaces for various distance threshold values in range between 10 – 500 nm at 40 nm step size. Contour levels = 0, 0.05, 0.25, 0.5, 0.75, 0.95 percentiles.

**Supp. Fig. 8.**
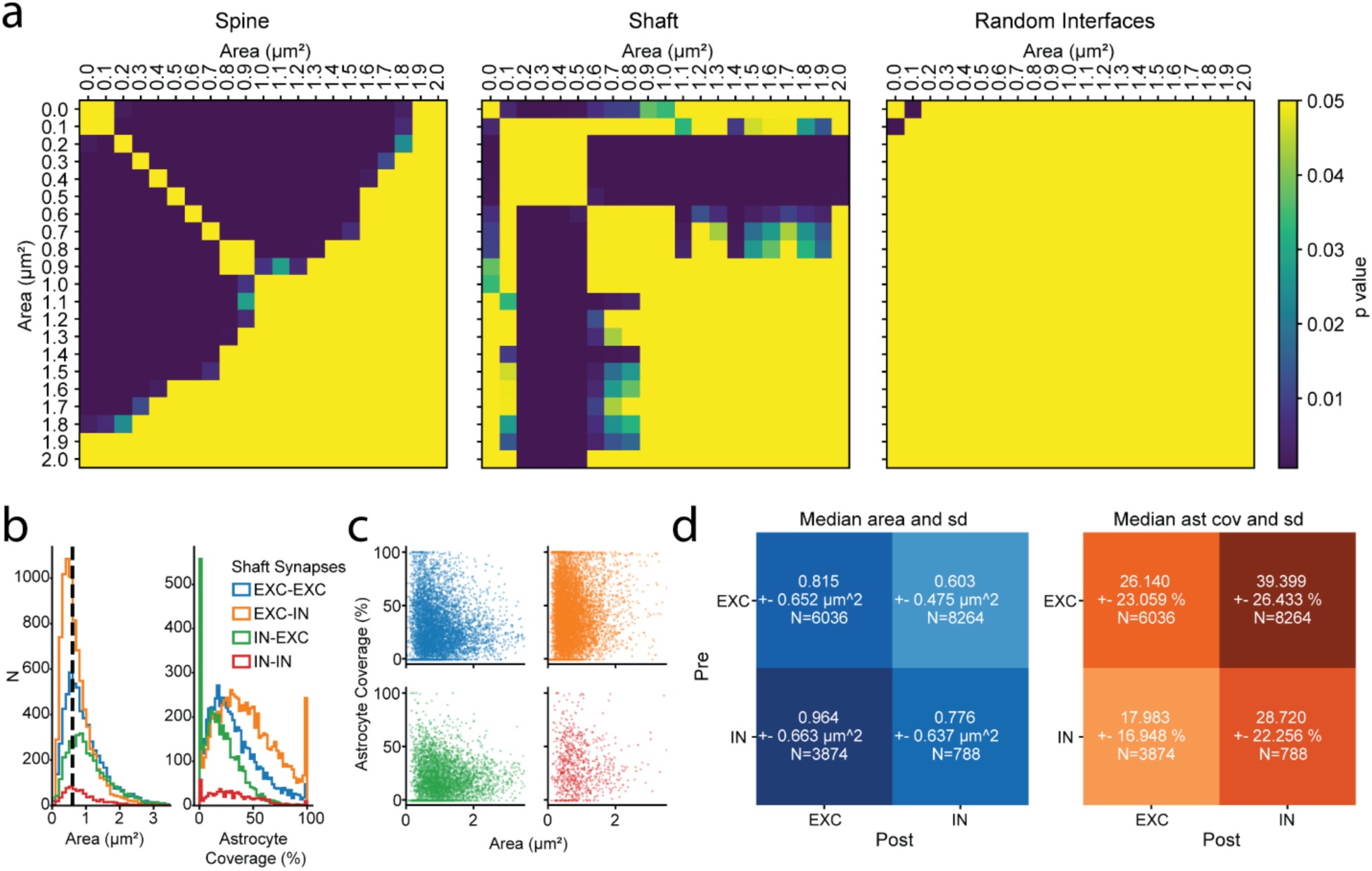
**a,** Post-hoc significance tests (Tukey’s HSD) for pairwise comparison of astrocyte coverage in different size bins, in reference to main figure panels Fig 2e and 3b. Darker color indicates significant differences in mean astrocyte coverage of each compared size bin. **b,** Distribution of shaft synapses’ contact area (left) and astrocyte coverage (right) for different pre- and post-synaptic neurite types (legend labeled as pre-to-post). Pre-synaptic EXC is an aggregation of CC and TC axons. Post-synaptic EXC is and aggregation of PD and AD dendrites. Post-synaptic IN is SD. **c,** Scatter plot of shaft synapses’ contact area vs astrocyte coverage, colors same as legend in b. **d,** Median, standard deviation, and sample numbers of shaft synapses’ contact area (left, ANOVA p<10^−297^, f: 476.41) and astrocyte coverage (right, ANOVA, p<10^−300^, f: 785.44) for different pre- and post-synaptic neurite types.

## Notes

### Competing Interest Statement

The authors have declared no competing interest.

